# Quantitation of Spatial Proteoforms in Alzheimer’s Disease

**DOI:** 10.64898/2026.06.30.735694

**Authors:** Daniel B. McClatchy, Natalie Turner, John R. Yates

## Abstract

Alzheimer’s disease (AD) pathogenesis involves complex, multifactorial changes to the brain proteome that conventional unfractionated analyses may obscure. Proteins frequently occupy multiple subcellular compartments as spatial proteoforms, yet the contribution of aberrant protein localization to AD pathogenesis remains poorly understood. To address this, we fractionated post-mortem human hippocampi from 13 AD and 14 non-AD individuals into four subcellular fractions and quantified 6,123 proteins by TMT-LC-MS. Although 75% of proteins were detected in more than one fraction, 78% of significant AD-associated alterations were restricted to a single fraction, demonstrating that subcellular localization is a primary determinant of disease vulnerability. Discordant abundance patterns between fractions revealed retromer complex mislocalization, nuclear transport dysfunction, and insoluble protein accumulation, with the endosomal-lysosomal and protein folding pathways most consistently perturbed. To examine how these perturbations evolve with disease progression, we applied the QUAD strategy to measure protein degradation in two fractions of APPswePS1delta9 mouse cortex at 2, 5, and 12 months. Degradation rates diverged between fractions and genotypes in an age-dependent manner, and cross-dataset comparison identified six proteins altered at the earliest pre-pathological timepoint, implicating vesicle transport and proteostasis disruption as initiating features of AD. These findings establish spatial proteoforms as essential units of pathogenic analysis and reveal disease-relevant signals invisible to bulk tissue approaches.

## Introduction

Alzheimer’s Disease (AD) is the most common form of dementia with an estimated 6.7 million people living with this disease in 2023 in the U.S and more than 55 million worldwide^1^. The number of people living with AD is projected to more than double by 2060. Caring for people afflicted with AD costs billions of dollars in the US yearly. AD is a neurodegenerative disease where signs of dementia begin at 65 years of age and older and then worsen until death. AD is definitively diagnosed only by post-mortem brain examination revealing two pathological hallmarks: extracellular amyloid plaques and intracellular neurofibrillary tangles (NFT). Plaques are composed of the amyloid beta (Abeta) peptide cleaved from the amyloid precursor protein (APP), and NFT are composed of the hyperphosphorylated Tau protein (i.e. MAPT gene). Arguably, Abeta has been the focus of drug therapy in the last decade, but recent approved drugs have been underwhelming with mild improvement and potential lethal side effects^2^. This does not exclude Abeta from pathogenesis; rather, Abeta-targeting drugs are ineffective at late disease stages. Thus, AD research is at a crossroads and new hypotheses are needed. We propose that effective AD drug development requires a holistic understanding of the complex, multifactorial AD brain proteome.

Proteomics via liquid chromatography coupled to mass spectrometry (LC-MS) is an excellent tool to dissect the complexity AD proteome. There have been phenomenal studies interrogating hundreds of AD brains with LC-MS providing novel insight into pathogenesis^3–5^. These reports quantified AD perturbations from unfractionated brain tissue. It has been estimated that more than 50% of all proteins are localized in more than one subcellular compartment^6^. Spatial proteoforms, defined as proteins with multiple subcellular localizations, may function differently depending on their compartment, as each environment provides access to distinct binding partners. Furthermore, the abundance of spatial proteoforms may differ between subcellular compartments. This can occur due to changes in protein transport, local translation, or local degradation. Biological fractionation of brain tissue grants the identification and quantification of spatial proteoforms^7^. We hypothesized that quantifying proteins across biological fractions would reveal novel aspects of AD pathogenesis, given that aberrant subcellular protein localization has been implicated in multiple human diseases. Translocation of the epidermal growth factor receptor to the nucleus can promote cancer^8^. Mutations in C9orf72 that cause the motor neurodegenerative disease amyotrophic lateral sclerosis (ALS) have been shown to disrupt nuclear-cytoplasmic transport (NCT)^9,10^. The RNA-binding protein TDP43 is depleted from the nucleus accumulates in the cytoplasm in ALS patients^11^. Mislocalization of APP within the endosomal network can alter its interactions with endopeptidases resulting in abnormal cleavage and increased generation of Abeta^12^. Disruptions in the vesicle transport is one of the earliest abnormalities in AD pathogenesis^13^. In addition, AD genetic risk factors include an abundance of proteins that regulate protein trafficking^14^. If vesicle transport is disrupted early in AD pathogenesis, this would suggest that protein mislocalization would be prominent in the AD proteome.

To understand the complexity of spatial proteoforms in pathogenesis, we fractionated AD and non-AD human hippocampi brain samples into four biological fractions. The hippocampus was chosen since it is essential to memory and vulnerable to AD pathology early in the disease progression. Each fraction was labeled with TMT isobaric tags and quantitated with LC-MS. The proteomes of the fractions were very similar, suggesting an abundance of spatial proteoforms. Most AD protein alterations, however, were limited to one fraction suggesting specific proteoforms are vulnerable to pathogenesis. Reciprocal alterations observed between the fractions suggest a portion of changes are due to protein transport. Since post-mortem brain tissue does not provide insight into early-stage AD perturbations, we also analyzed an AD mouse model. We quantitated protein degradation in different biological fractions in the APPswePS1delta9^15^ AD mice at 2, 5, and 12 months. Overall, this study reveals vulnerabilities of specific spatial proteoforms in AD pathogenesis and demonstrates that protein degradation is altered by Abeta pathology.

## Material and Methods

### Human brain samples

Fresh frozen human postmortem hippocampi from neuropathologically confirmed AD and cognitively normal control cases was obtained from the Shiley-Marcos Alzheimer’s Disease Research Center of the University of California, San Diego.

### Mice

Transgenic Alzheimer’s disease hemizygote mice (B6.C3-Tg (APPswe,PSEN1dE9)85Dbo/Mmjax; #034829-JAX)^16^ were purchased from the Jackson laboratories and bred with C57BL/6 mice about from the Scripps Research breeding colony. All animals were housed in plastic cages located in a temperature- and humidity-controlled animal colony and were maintained on a reversed day/night cycle (lights on from 7:00 P.M. to 7:00 A.M.). Animal facilities were AAALAC-approved, and protocols were in accordance with the IACUC. Inotiv (West Lafayette, Indiana) prepared AHA pellets as previously described^17^. Both females and male mice were used for this study. AD mice and their WT littermates were given a standard mouse diet until the ages of 2, 5, or 12 months. At these ages, mice were given an AHA diet ad libitum for 4 days, then returned to normal diets for 7 days. Mice were sacrificed by isoflurane inhalation after four days on the AHA diet or 7 days of the standard diet post the AHA diet. The brains were quickly dissected and snap-frozen in liquid nitrogen.

### Abeta Elisa

The human samples were in 4mM Hepes, 200mM NaCl at a concentration of 1mg/ml. Samples were centrifuged for 10min at 21,000 x g. The supernant was analyzed with the Abeta 1-42 Human Ultrasensitive Elisa (**#** KHB3544;ThermoFisher, Carlsbad, CA) following the manufaacturer’s protocol. For mice, 1mg of cortical homogenate was resuspend in a total of volume 100ul using PBS. The samples were sonicated at 20% amplitude for 10 sec. twice using a Qsonica(Newtown, CT) model#125 tip sonicator. The 2month samples were analyzed undiluted, and the 5month and 12month samples were diluted 1:4 and 1:100, respectively, with supplied diluent. Wild-type brain samples were analyzed at each timepoint but a signal was not observed.

### Brain fractionation

Each human hippocampi was homogenized on ice in a Teflon dounce grinder with cold 4mM HEPES, pH7, 200mM NaCl (H1 buffer) with phosphatase and protease inhibitors(Pierce). Protein concentration was determined by a BCA assay (Pierce). Two milligrams of the brain homogenate in 2ml of 4mM HEPES, pH7, 200mM NaCl was centrifuged at 700 x g for 10min. All centrifugation steps were performed at 4°C. The supernant (S1) and the pellet (P1) were separated and frozen. S1 was centrifuged at 17.2K x g for 10min. The supernant (S2) and the pellet (P2) were separated and saved for further processing. H1 buffer with 0.5% NP-40 was added to the P1 pellet and incubated at 4°C while shaking overnight. Then, the P1 pellet was centrifuged at 700 x g for 10min. The supernant (S3) and the pellet (P3) were separated and frozen. After thawing, S2, P2, S3, and P fractions were lysed with 0.5% SDC, 0.5% SDS in H1 buffer. Samples were incubated at 37°C while shaker, then tip sonicated (Qsonica) at 50% amplitude for 10sec three times. Protein concentration was determined by a BCA assay (Pierce). For mice, cortices were homogenized in a teflon dounce grinder on ice in PBS with protease and phosphatase inhibitors (Roche, Indianapolis, Indiana). After homogenization, protein concentration was determined with a Pierce BCA protein assay (Life Technologies, Grand Island, NY). Then, tissue was fractionated using the same protocol as the human brain fraction. Only S2 and P3 fractions were prepared for further analysis. For each timepoint, the Day0 samples for each genotype with an equal number of male and female mice were pooled for each fraction. Each biological replicate(N=4) for the Day7 samples were pooled from one male and one female mouse cortex.

### Click Chemistry

Sixteen click reactions were performed for each fraction at each timepoint using 2mg of fractionated cortical tissue for each reaction for the following conditions: AD-Day0, WT-Day0, AD-Day7, and WT-Day7. Prior to the click reactions, the samples were precipitated with 4X the volume of cold −20°C acetone. The dried precipitated pellets were resuspended with 0.025% SDS in PBS. The click reactions were performed as previously described with the biotin-alkyne DADPS^18^. The click reactions were incubated for 1hour on a 30°C thermomixer, the overnight at 4°C while rotating.

### Protein Digestion and TMT labeling

For human samples, twenty-five micrograms of protein sample was precipitated with methanol and chloroform. The dried precipitated pellet was resuspended with 8Murea in 100mM Hepes, pH 8.5. The protein was reduced with 5mM TCEP at 55°C for 20min, then alkylated with 10mM chloro-iodoacetamide at room temperature in the dark for 20min. The urea was diluted to 2M with 100mM Hepes, pH 8.5 and 1ug of trypsin was added. The digestion was performed on a 37°C shaker overnight. The digestions were desalted, then dried with a speed-vac. The dried peptides were resuspended in 10ul of 0.5M Hepes, pH 8.5. Two microliters from all peptide samples were pooled and used reference channels. TMT 16plex 0.5mg vials were resuspended in 20ul of acetonitrile. One microliter (i.e. 25ug) of TMT was added to the peptides and incubated for 30min, then the incubation was repeated with another 25ug. After TMT labeling efficiency was confirmed by LC-MS, 2ul of 5% hydroxylamine was added to each of the peptide samples. For each fraction, the peptide samples were combined into two 16plex sets each with a reference channel. Each TMT set was fractionated with Pierce™ High pH Reversed-Phase Peptide Fractionation Kit following the manufacturer’s instructions except the wash was increased to 10% acetonitrile and the last fraction contained 80% acetonitrile. For the mouse samples, the click reactions were precipitated, then digested with trypsin as previously described above. Each click reaction was incubated with 100µl of Neutravidin agarose (Thermo # 29200) overnight at 4°C while rotating. After washing with PBS, the AHA peptides were eluted twice with 5% formic acid. The elutions were then desalted and labeled with 16-plex TMT and fractionated as described for the human samples.

### LC-MS

The TMT labeled samples were analyzed on an Orbitrap Eclipse Tribrid mass spectrometer (Thermo). Samples were injected directly onto a 25 cm, 100 μm ID column packed with BEH 1.7 μm C18 resin (Waters). Samples were separated at a flow rate of 300 nL/min on an EasynLC 1200 (Thermo). Buffer A and B were 0.1% formic acid in water and 90% acetonitrile, respectively. A gradient of 1–25% B over 120 min, an increase to 40% B over 40 min, an increase to 100% B over 10 min and held at 100% B for a 10 min was used for a 180 min total run time. Peptides were eluted directly from the tip of the column and nanosprayed directly into the mass spectrometer by application of 2.5 kV voltage at the back of the column. The Eclipse was operated in a data dependent mode. Full MS1 scans were collected in the Orbitrap at 120k resolution. The cycle time was set to 3 s, and within this 3 s the most abundant ions per scan were selected for CID MS/MS in the ion trap. MS3 analysis with multinotch isolation (SPS3) was utilized for detection of TMT reporter ions at 60k resolution. Monoisotopic precursor selection was enabled and dynamic exclusion was used with exclusion duration of 10 s.

### Bioinformatics

ProteomeDiscoverer 2.5 was used to analyze the MS/MS data to identify and quantified peptides. The MS spectra was searched using the Uniprot mouse protein database with isoforms(version v2024-01-24) or Uniprot human protein database with isoforms(version v2022-08-03) and a common contaminant proteins list. The decoy database was the reverse of this Uniprot database to filter identifications to a 1% FDR. The peptides were allowed to have a maximum of two miscleavages with a minimum of 7AA. The static modifications searched for were TMT on lysine and peptide N-terminal (+ 304.2071Da) and cysteine carbamidomethylation (+57.021464 Da) for the human analysis. For the mouse analysis, an additional differential modification on methionine (79.0711) was searched for the DADPS modification on AHA. Reporter ion distributions specific to the lot number of the TMT reagent were employed as correction factors. For the AHA analysis, only AHA containing peptides were quantified and further analyzed. For the human analysis, ANOVA was performed in ProteomeDiscoverer 2.5. For the mouse analysis, the quantified proteins were further processed in Excel. In each of datasets, there were 4 WT Day0 measurements, 4 AD Day0 measurements, 4 WT Day7 measurements, and 4 WT Day7 measurements. For each genotype, each Day7 protein TMT measurement was divided by the median Day0 protein TMT value to calculate the extent of protein degradation or the Day7/Day0 ratio. Student t-test determined the significance protein differences between Day7/Day0 WT ratios and Day7/Day0 AD ratios. The ratios were log^2^ transformed prior to the statistical analysis.

WGCNA was performed by MetaNetwork^19^. The input was the normalized protein TMT intensities from PD. Proteins with missing values were removed from the analysis. The mean module eigenprotein values that were significantly different (p < 0.05) were analyzed by Webgesalt to determine GO biological processes (BP) that were significantly enrichment ( using of 5% FDR filter^20^. The top three BP for each significant module was used for further analysis. Some modules had less than three BP that passed the filters. For some modules, BP were combined into one if it was determined that a subset BP with >50% proteins identical and then the BP with the most proteins was reported. PACOM was used to the uniqueness of the protein identified in the human fractions^21^. Webgestalt performed the GO enrichment analyses^22^. Uniprot was used to annotated mitochondrial proteins. Prism Graphpad and MicrosoftExcel were used for graphing and additional statistical analyses. Cytoscape were used for visualization of protein networks^23^.

### Immunoblot Analysis

Samples were solubilized with 4X Laemmli Sample Buffer (Bio-Rad) with β-mercaptoethanol, separated with 4–12% Bis-Tris gradient gel(Life Technologies), transferred to PVDF blotting paper, and developed as previously described^24^. The immunoblotting antibodies were VPS-35(SCBT, #sc-374372), TAU-5(Life Technologies, #AHB0042), and Ubiquitin (CST, #431224). AcquaStain Protein Gel Stain (Bulldog-Bio) was used as a total protein stain on gels. The immunoblots were quantitated as previously described^7^.

## Results

Twenty-seven post-mortem human hippocampi samples were analyzed. Thirteen samples were from AD patients and fourteen were from age-matched non-AD (NA) individuals (*Table S1*). The NA samples were comprised of two different classifications: 1) humans (N=11) with AD brain pathology (i.e. Tau and/or Abeta), which is also known as asymptomatic AD (AsymAD) or 2) humans (N=3, i.e. “normal”) with no AD brain pathology. An Abeta^42^ elisa was performed on total brain homogenate, which confirmed the post-mortem classifications *(Figure S1A).* The samples were processed as depicted in Figure 1. Each sample was fractionated with differential centrifugation resulting in four fractions: S2, P2, S3, and P3. The fractions were then digested with trypsin and labeled with TMT isobaric tags to quantitate the differences between AD and NA in each fraction using LC-MS. There were 6123 unique proteins identified with at least two peptides among all the fractions (*Table S2*). A similar number of proteins were identified within each fraction, and 75% of the identified proteins were observed in more than one fraction (Figure 2A and B). The P3 fraction contained the most unique proteins (Figure 2C). We analyzed the unique proteins identified in each fraction to provide insight into distribution of cellular compartments using Gene Ontology (GO) (Figure 2D and *TableS3*). P3 was significantly enriched in complexes associated with the nucleus, whereas P2 contained mitochondrial components. S2 and S3 fractions had more diverse subcellular compartments significantly enriched. Lysosomes and vesicles were enriched in the S2 fraction, whereas the S3 fraction was enriched for the endoplasmic reticulum (ER).

**Figure 1.**
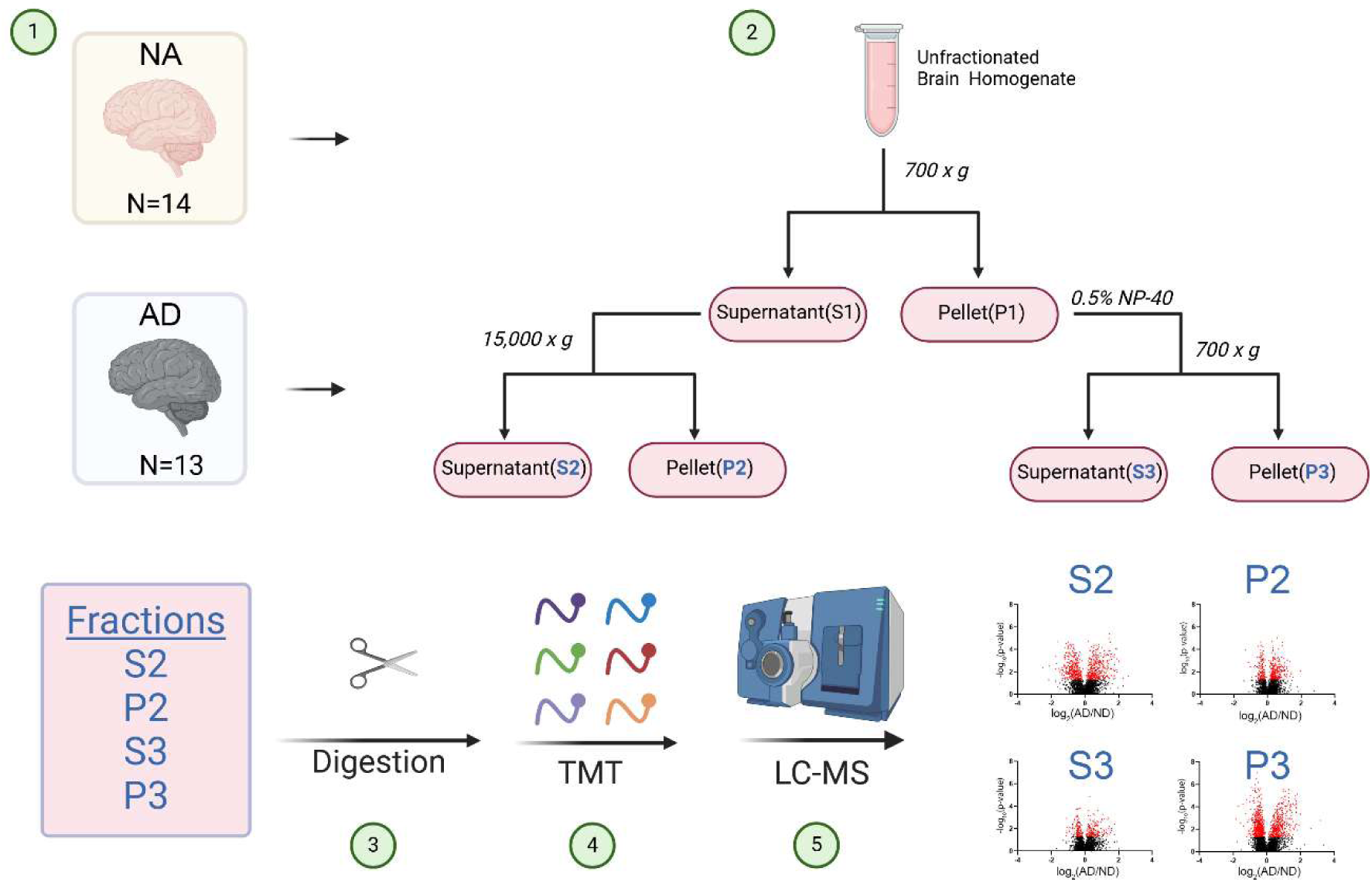
Schematic of the human brain analysis. Post-mortem hippocampi from fourteen non-demented (ND) and thirteen Alzheimer’s disease (AD) diagnosed individuals were employed in this study. Each hippocampus was individually fractionated to generate four fractions: S2, P2, S3, and P3. Each fraction was digested with trypsin, and then the resulting peptides were labeled with isobaric TMT tags. Labeled peptides were combined to generate four multiplex samples (i.e. S2, P2, S3, and P3) for LC-MS analysis. The output was the statistical differences between AD and ND in each fraction.

**Figure 2.**
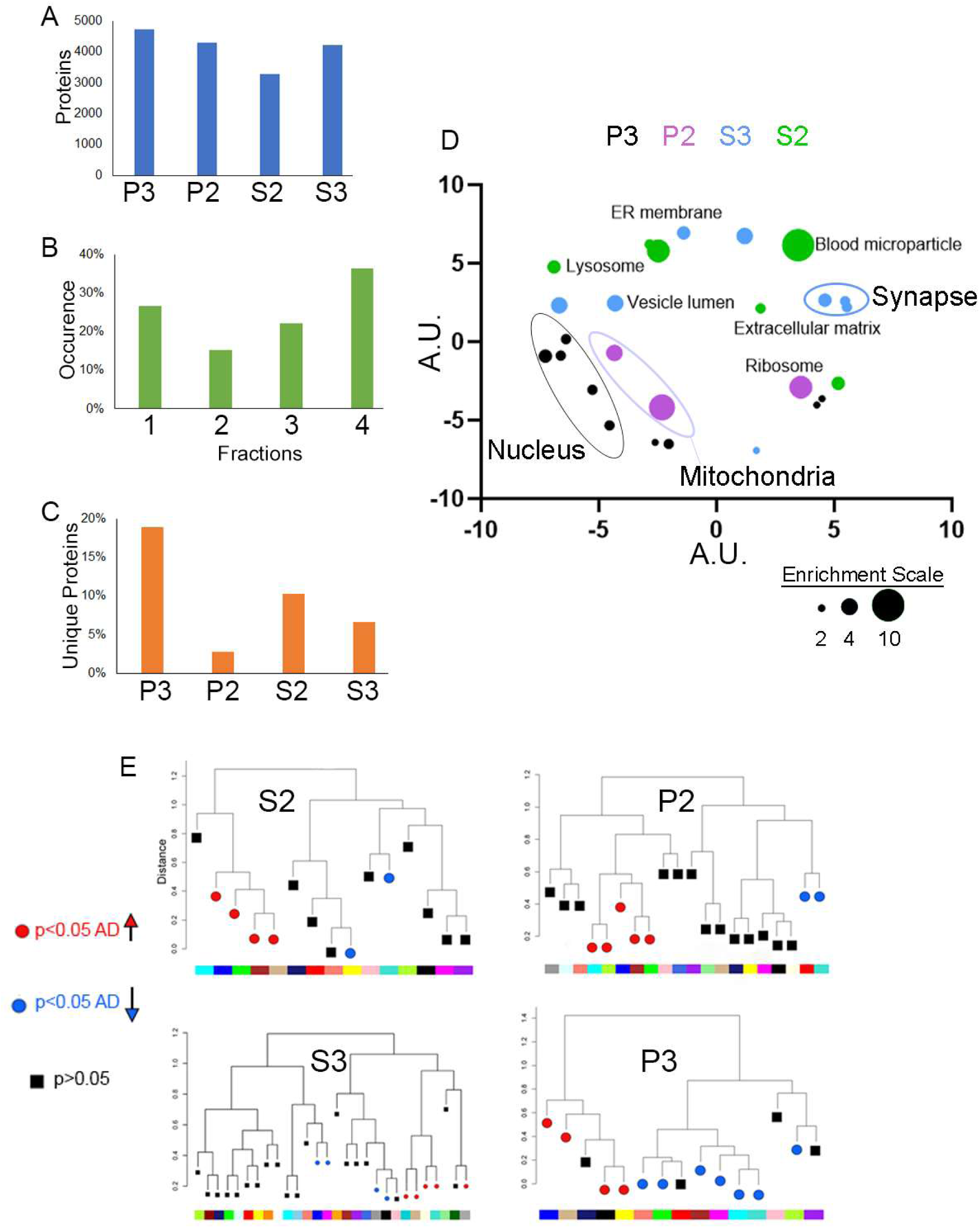
**A**, The number of proteins identified in each fraction with at least two peptides. **B**, The percentage of proteins identified in one or more fractions. **C**, The percentage of unique proteins identified in each fraction. **D**, Enriched GO terms (colored circles) from the unique identifications from each fraction were clustered for similarity using the algorithm REVIGO. Colors represent each fraction: P3 (black), P2 (purple), S3 (blue), and S2(green). GO terms from the same subcellular compartment (i.e. nucleus) were labeled. Size of each circle represents the enrichment of the GO term. Numbers on the axes are arbitrary units (A.U.) **E**, Module eigenproteins (MEs) generated by WGNCA for each fraction. Each ME was assigned an identifying color as indicated by the color strip below each hierarchical fraction map. Red and blue MEs indicate significantly (p < 0.05) up- and down-regulated in AD, respectively. Black MEs were not significant (p > 0.05).

### WGCNA of human brain fractions

We performed WGCNA (Weighted Gene Correlation Network Analysis), which is an unsupervised construction of cluster-based modules, to extract biological insight from the TMT quantified proteins^19^. Each module is represented by an eigenprotein; the mean module eigenprotein (ME) expression is then used to identify significant differences between conditions. This was performed for each fraction. We identified 16 MEs for both S2 and P3 and 21 and 25 for P2 and S3, respectively (Figure 2E and Supplementary Table4). P3 had the highest percentage (69%) of significantly different modules (Figure S1B). We identified the enriched GO biological pathways from the MEs that were significantly altered in AD (Figure 3A and Supplementary Table5). Vesicle and protein trafficking pathways were prominently altered, including vesicle-mediated transport at synapses and between endosomal compartments, cytoskeleton-dependent intracellular transport, and protein localization to the cell periphery, plasma membrane, and ER. There were six pathways that were regulated differently between fractions: generation of energy, translation initiation, protein folding, protein localization to ER, mitochondrial respiratory chain complex assembly, and nucleoside monophosphate metabolism (Figure S1C). Most proteins annotated to these pathways were observed in only one fraction (Figure S2). There were, however, some shared proteins between fractions even though they were observed in modules that had an opposite ME expression pattern. For example, protein folding pathway was upregulated in the blue and green modules in the S2 with AD but downregulated in the turquoise modules in the P2 and P3 fractions (Figure 3B). There were 86 proteins that were annotated to protein folding in these fractions. Fourteen proteins were shared between two fractions, and four proteins were quantified in all three fractions (Figure 3C). Of particular interest, CCT1 was quantified in all three fractions and is a component of an eight subunit (CCT1-8) chaperonin complex. In addition, the subunit PFDN2 of the prefoldin chaperone complex, which binds the CCT chaperonin^25^, was also quantified in all three fractions. Other chaperonin and prefoldin complex subunits were quantified, but most were unique to one fraction (Figure 3D). For example, CCT5 and CCT8 were quantified in P2 and CCT2 and CCT3 were observed in S2. This corresponds to emerging evidence of non-canonical CCT chaperonin structures^26^ and indicates they are differentially regulated in AD. Overall, the WGCNA demonstrates that perturbations in the AD brain proteome can be separated into different biological fractions.

**Figure 3.**
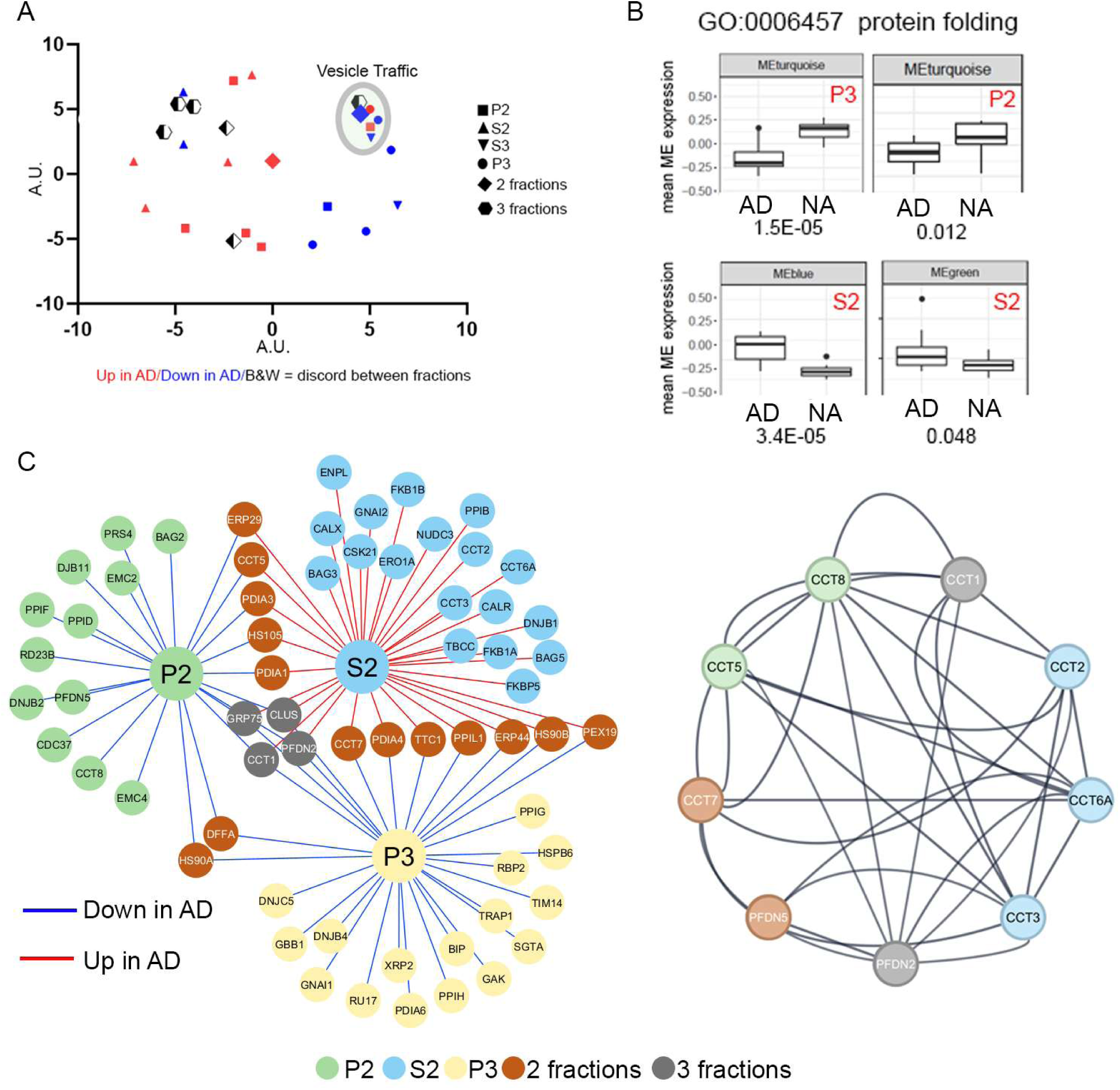
**A**, Enriched GO biological processes in significantly different MEs shown in Figure 2E. GO terms were clustered for similarity using the algorithm REVIGO. Symbol shape represents the fractions the enriched GO term was observed. Colors represent if the GO term was enriched in a ME that up-regulated (red), down-regulated (blue), or both up- and down-regulated (black and white) in multiple fractions with respect to AD. GO terms within the green circle are related to vesicle and protein trafficking. **B**, Mean expression for MEs that were enriched for the GO term protein folding. The p-value for each AD vs ND comparison is below each graph. **C**, *Left panel*: Proteins annotated as protein folding observed in B. Colored nodes represent if proteins were quantified in P3 (yellow), P2(green), S2(blue), or multiple fractions (2 fractions-brown, 3 fractions-grey). Edge color indicates that the protein assigned to a ME that was up-regulated (red) or down-regulated (blue) in AD. *Right panel*: Proteins in the left panel that are subunits of the CCT chaperonin and prefoldin complexes. Edges represented physical interactions assigned by STRING database with 0.9 confidence.

### ANOVA of human brain fractions

Next, we performed a complementary statistical analysis (i.e. ANOVA) on each fraction, and the highest number of significant changes were observed in the P3 and S2 fractions (Figure 4A, B and Supplementary Table6). There were 2405 statistical changes between AD and NA when all fractions were combined, which encompassed 1905 unique proteins. Interestingly, 78% of these significant changes were specific to one fraction even though 89% of these proteins were quantified in more than one fraction (Figure 4C). A meta-analysis of seven proteomic TMT datasets on human unfractionated AD non-hippocampal brain tissue confirmed 748 (31%) AD changes in our dataset (Supplementary Table 7)^27^. Enrichment analysis on significant changes from all the fractions revealed an enrichment in GO biological processes connected to the vesicle and protein trafficking including autophagy, vesicle transport, Golgi-mediated transport, and protein localization to membranes (Figure 4D and Supplementary Table 8). Inflammation and energy production were also enriched and previous implicated in AD pathogenesis^28^. Among the observed changes, there were 11 AD risk factors: ANK3, APOE, APP, BIN1, CTSB, CLU,COX7C, GRN, FAK, FERMT, and SNX1 (Figure 4E). All risk factors were altered in one fraction, except for Bin1 which was downregulated in both S2 and P2. We further investigated the peptides assigned to APP, which were significantly altered only in P3. We separated the quantified APP peptides that were within the Abeta using the Abeta1-42 sequence and outside this sequence (Figure 4F). Although LC-MS analysis cannot determine if an identified Abeta peptide was cleaved in vivo from APP or cleaved in vitro with trypsin, the most Abeta peptides were quantified in the P3 fraction. Abeta peptides were quantified in all fractions, however, and up regulated in all the fractions in AD. APP peptides not including the Abeta sequence were also quantified in all fractions, but there was no difference between AD and NA. These non-Abeta peptides were the least frequent in P3 among the fractions.

**Figure 4.**
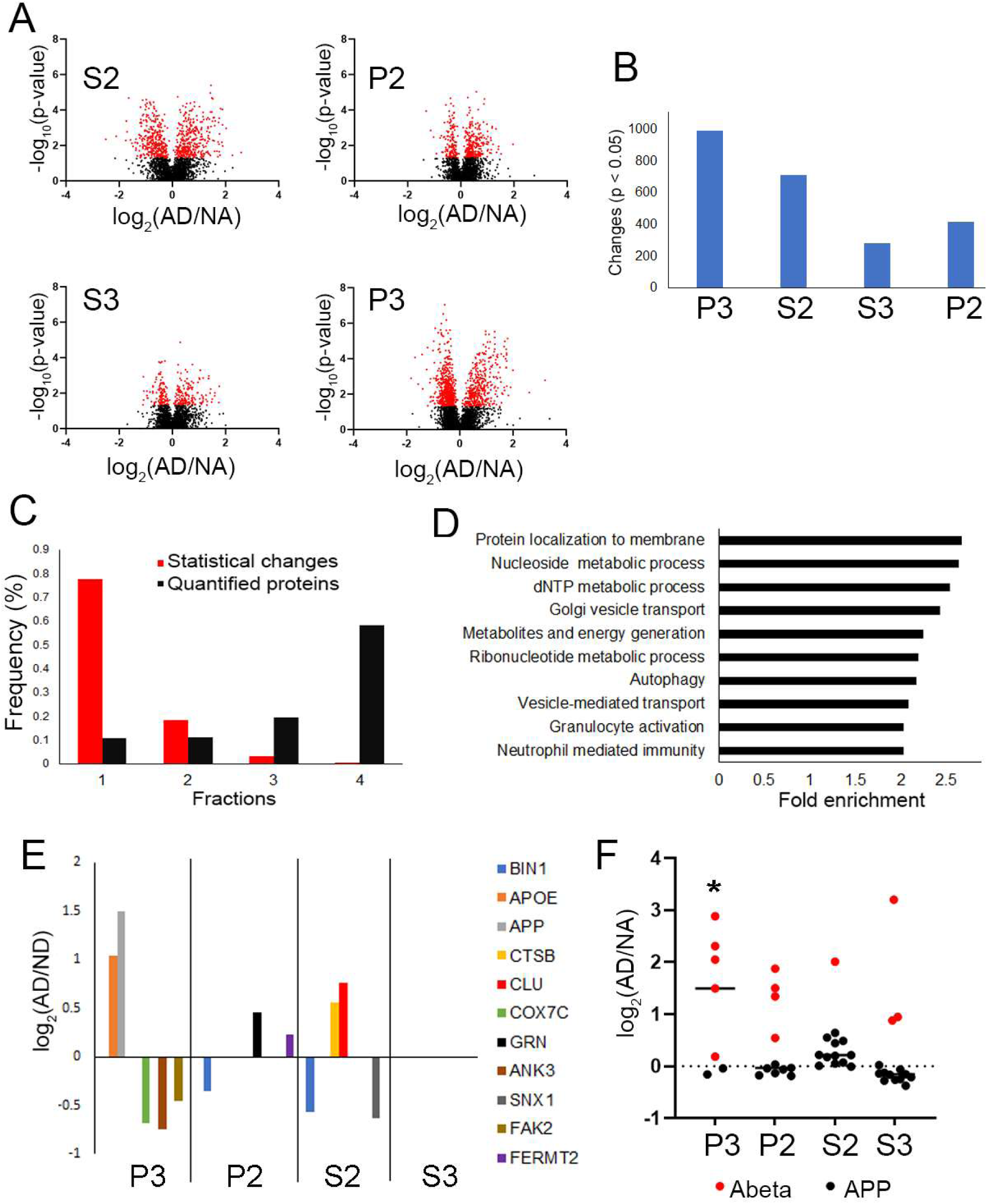
**A**, Volcano plots of the ANOVA of each fraction. Red circles indicate the proteins with a p-value < 0.05 whereas black circles indicate a p-value < 0.05. X-axis is the log_2_ of AD/ND protein rations. **B**, The number of significant changes plotted for each fraction as shown in A. **C**, GO biological processes significantly enriched from the proteins in B. X-axis is -log_10_ of the pvalue of the significance of the enrichment. Red line represents p-value 0.01.D, The frequency of the changes (red) in B in the fractions and the frequency of the proteins in B in the fractions that were quantified (black). E, AD genetic risk factors that were observed significantly altered in A. F, Quantified peptides from the Abeta sequence (red) and APP peptides without the Abeta sequence (black). Y-axis is log_2_ transformation of AD/ND peptide ratio. * p value < 0.05.

We also examined in more detail the protein Tau. Tau was significantly increased in AD in P3 and S3 (Figure 5A). We confirmed our LC-MS quantification with immunoblots using a Tau antibody (Figure 5B and FigureS3A and B). The immunoblots revealed robust detection of high molecular weight Tau (HMWtau) in all the AD fractions but little to none in the NA fractions. HMWtau can form aggregates and promote Tau propagation^29–31^. Next, we performed correlation analysis between Tau abundance and the Braak score in each fraction. Tau propagates into anatomically defined regions corresponds to the severity of AD. This sequential spreading of Tau is assessed at autopsy by probing brain sections with a tau antibody that recognizes aggregated phosphorylated Tau using the Braak scale^32^. The scale ranges from 1 to 6 and 6 is the most severe. We observed a significant correlation between Braak scores and Tau abundance in P3 (r= 0.67, p < 0.0002) and S3 (r= 0.46, p < 0.02) AD fractions (Figure 5C). Furthermore, we hypothesized that proteins correlating with Tau in AD samples may serve as biomarkers or contributors to Tau pathology, whereas those correlating with Tau in NA samples may reflect Tau resilience. Many studies have been performed to identify proteins that regulate Tau expression and aggregation^33–37^. To identify potential Tau regulators, we identified proteins that were correlated (r >0.7) with Tau abundance in each fraction in the AD and NA samples. We identified 124 proteins that correlated with Tau and 15 proteins were previously reported as a Tau regulator (Supplementary Table 9)^34,35,38^. Most proteins were unique to one fraction and only 13 proteins were observed in more than one (Figure 5D). The autophagy regulator, Rab2a was correlated with Tau in the P2, S3, and S2 fractions in only NA samples suggesting that it may be a negative regulator of Tau aggregation. Rab2a was identified as a regulator of Tau oligomers in genome-wide CRISPRi screen in induced pluripotent stem cell (iPSC)-derived neurons ^38^. A literature search reveals many of these correlated Tau proteins have previously been associated with the pathogenesis of AD or other dementias. For example, FERMT2 was correlated with Tau in AD-P3 and is a genetic AD risk factor for AD^39^. Modulating FERMT2 expression can regulate Tau pathology^40^. Overall, our ANOVA demonstrates that even though a protein is identified in multiple fractions, most of the disease related changes are localized in one fraction. Furthermore, the P3 fraction is significantly enriched in proteins (i.e. Abeta and Tau) prone to form insoluble aggregates in AD.

**Figure 5.**
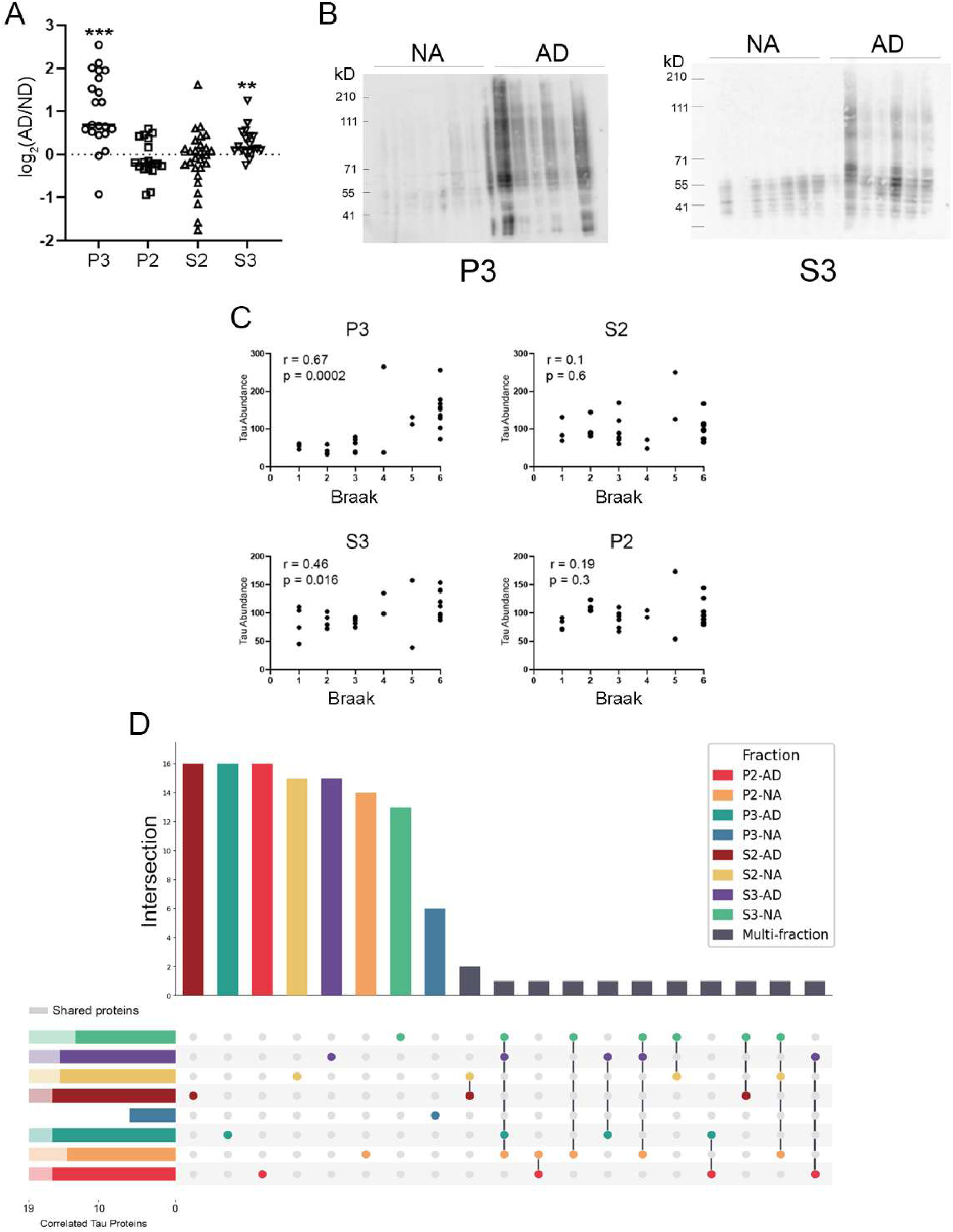
**A**, Tau peptides quantified in each fraction. Y-axis is log_2_ transformation of AD/NA peptide ratio. **B**, Immunoblots were probed with Tau antibody for P3 and S3 fractions. The Tau immunoreactivity is graphed on the right plot after being normalized to a total protein gel stain. The y-axis is normalized pixel intensity (arbitrary units). **C**, The average Tau protein abundance (y-axis) quantified by LC-MS in each human hippocampal sample correlated to its Braak score (x-axis) for each fraction. The correlation factor (r) and the p-value of the correlation is listed in the upper left of graph. **D**, The overlap of proteins that correlated with Tau abundance in each fraction. The bars in the upper graph represent the number of unique proteins.

### Gene expression changes in Human brain fractions

If protein abundance is altered by transcription, the perturbation would be expected to be observed in all fractions where the protein is quantified. Fifty-five percent of the significantly altered proteins had the same direction of change in more than one fraction, which may represent disease alterations generated gene expression (Figure 6A). Eight proteins were upregulated in AD across all fractions; none were consistently downregulated in all fractions (Figure 6B and Figure S3C-E). These consistently upregulated proteins are involved in regulation of the cytoskeleton and immune response. The P3 and P2 fractions shared 102 protein expression patterns, which was the highest agreement among all the possible fraction comparisons (Figure 6C). These P3/P2 proteins were enriched in multiple biological processes including cell adhesion processes, receptor mediated endocytosis, platelet degranulation, and cation transport (Figure 6D and Supplementary Table 10). The proteins annotated to these pathways were upregulated in AD except for cation transport, where the proteins were downregulated. Three proteins (ATP6V1A, ATP6V1B2, ATP6VG2) within this category are subunits of the V1 complex which comprises the larger 31 subunit vacuolar(H+)-ATPase (V-ATPase) (Figure 6E and Figure S3D). V-ATPase is essential for neurotransmission and the acidification of lysosomes and endosomes to support vesicle trafficking and protein degradation^41,42^. The complex has been observed to be dysfunctional in AD and proposed as a therapeutic AD target by multiple laboratories^5,43^.

**Figure 6.**
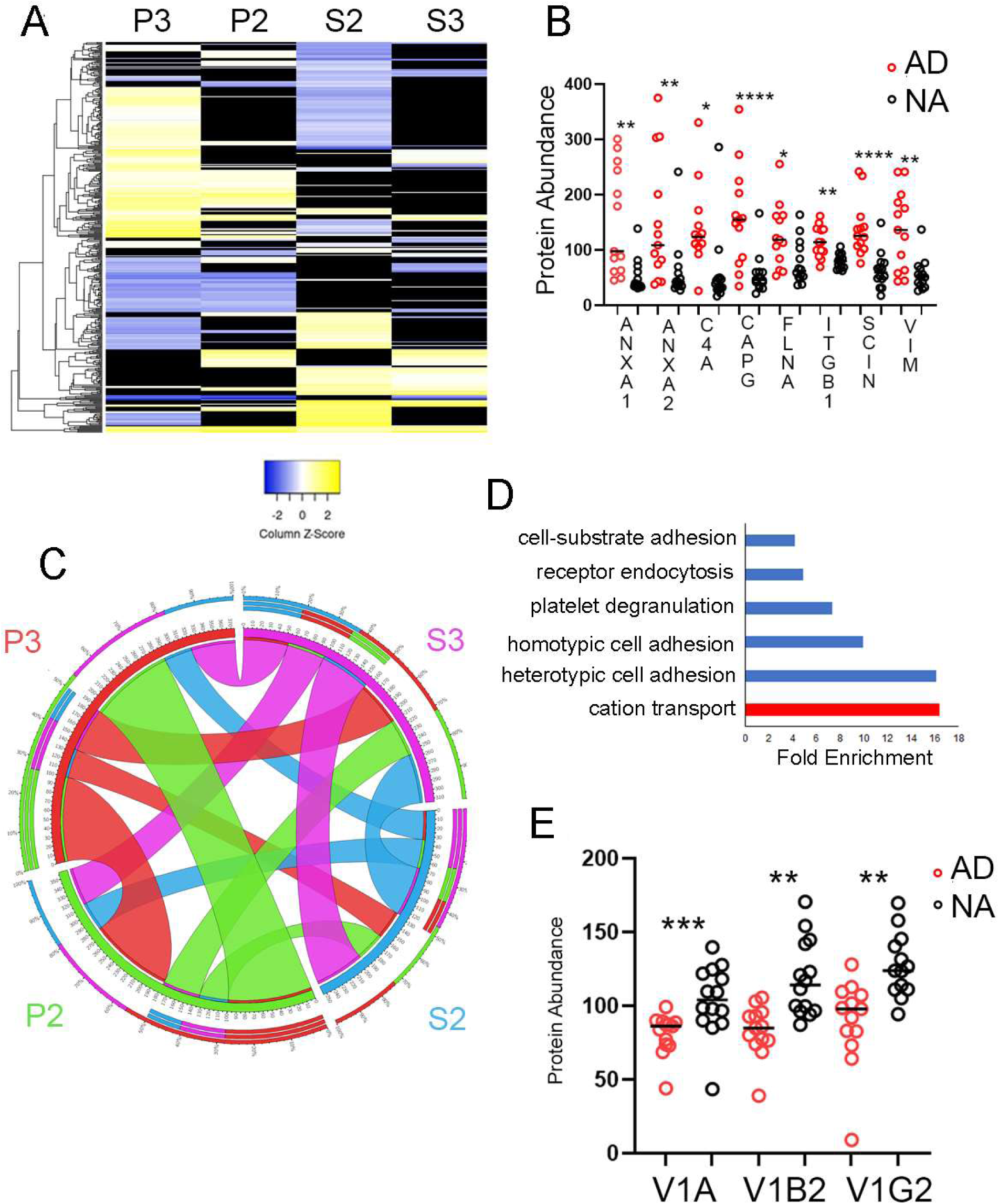
**A**, Heatmap of all the significant ANOVA protein changes observed. Y-axis is proteins. Yellow indicates up-regulation in AD, blue indicates down-regulation, and black indicates no significant change. **B**, Proteins that were observed to be altered in all four fractions. Graph shows the TMT protein quantification values for the AD (red) and ND (black) samples from the P3 fraction analysis. Y-axis is TMT protein abundance (arbitrary units). **C**, Circos plot of significant proteins where the abundance alteration in AD hippocampi samples concurred between fractions. **D**, The significant enriched GO biological processes from the shared proteins from the P2 and P3 fraction analyses. Blue bars indicate up-regulation in AD and red represents down-regulation in AD. **E**, The TMT protein quantification values for proteins (ATP6V1A, ATP6V1B2, ATP6VG2) of the V-ATPase complex in the P2 fraction. Each circle represents either an AD (red) or ND (black) sample. Y-axis is TMT protein abundance (arbitrary units). * < 0.05, ** < 0.01, *** < 0.001, **** < 0.0001.

### Translational changes in human brain fractions

Next, we examined proteins showing opposite expression patterns between fractions (Figure 7A). We observed the largest number (i.e. 150 proteins) of shared proteins with discord measurements between the P3 and S2 fractions. Sixty-six percent of these proteins were increased in P3 in AD and decreased in S2 in AD, whereas the other 34% had the opposite trend. Among these discord proteins, the most enriched protein functional group with a 10-fold enrichment was Ran GTPase binding (Supplementary Table 11). Ran GTPase is essential for nuclear-cytoplasmic transport (NCT). All the Ran GTPase binding proteins had the same pattern of an increase in the AD P3 fraction and decreased in the AD S2 fraction (Figure 7B). These quantified binding proteins are required for either nuclear import (KPNB1, RANBP7, RANBP5) or export (RANBP16). RANBP5 is required for nuclear import of RPL23 which was downregulated in AD P3^44^. With the enrichment of nuclear proteins in P3, this suggests inefficient nuclear transport in AD, which has been previously reported^45^. This indicates that other nuclear proteins may be mislocalized in our dataset. Nuclear proteins accounted for 36% of these discord P3/S2 proteins. These nuclear proteins formed a functional network with an enrichment (p value 2.44x 10-05, FDR 0.005) of the spliceosome complex (Figure 7C), which has also been reported to be dysregulated in AD^46^. In addition to Ran GTPase binding, multiple other protein functional groups were enriched that were related to the cellular response to misfolded proteins including ubiquitin binding activity, ubiquitin ligase activity, protein folding, and heat shock binding activity (Supplementary Table 11). These protein functional groups may be localized to the AD P3 fraction due to an enrichment of misfolded or aggregated proteins. This is consistent with the significant increase of known misfolded and aggregated proteins (i.e., Abeta and Tau) in AD P3. Another non-nuclear protein that was enriched in AD P3 was VPS35, which is one of the core subunits of the retromer complex. The retromer complex transport proteins from the endosomes to the TGN and the plasma membrane. Disruption of VPS35 function can disrupt vesicle transport leading to accumulation of misfolded proteins and has been hypothesized to contribute to AD pathology^47,48^. VPS35 was upregulated in the AD P3 fraction and downregulated in the AD S2 fraction, and we observed the same expression pattern for its binding partner, VPS29 (Figure S3F and G). This aberrant VPS35 AD expression pattern between fractions was confirmed by immunoblot analysis (Figure 7D).

**Figure 7.**
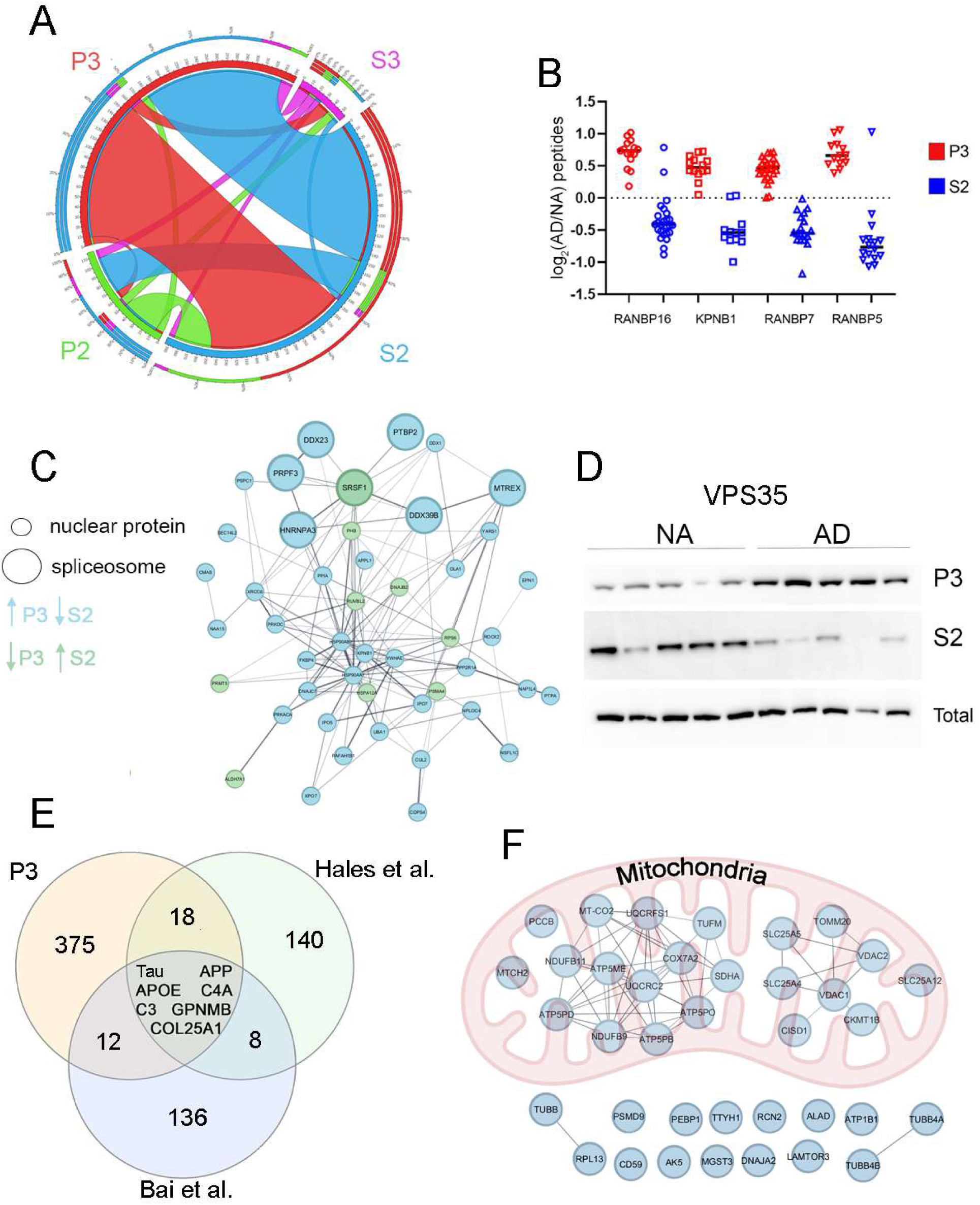
**A**, Circos plot of significant proteins where the abundance alteration in AD hippocampi samples disagreed between fractions. GO molecular functions significantly (p < 7.7 x 10^-4^, FDR < 0.05) enriched in discord P3/S2 proteins. **B**, The quantified peptides for RAN GTPase proteins that were up-regulated in the P3 fraction (red) and down-regulated in the S2 fraction (blue). The y-axis is the log_2_ of ratio of the AD peptide measurement over the ND peptide measurement. **C**, Network of the nuclear proteins from the P3/S2 discord dataset generated from String algorithm. Large circles denote proteins that are part of the spliceosome complex. Blue nodes denote proteins up-regulated in P3 and down-regulated in S2, whereas green nodes are proteins with the reverse expression pattern. **D**, Immunoblots analyzing S2, P3, and unfractionated (total) human hippocampi samples with a VPS35 antibody. **E**, The overlap of proteins quantitated to be up-regulated in the AD P3 fraction to proteins from two published studies (Bai et al. 2013 and Hales et al. 2016) that were quantitated to be up-regulated in the AD insoluble fraction. **F**, Proteins that were significantly decreased in P3 AD and significantly increased in S2 AD. More than half of the proteins are localized to the mitochondria. Edges denote an interaction using String database confidence 0.7. p-values: * < 0.05, ** < 0.01, ***< 0.005, **** < 0.0001.

These non-nuclear proteins upregulated in AD P3 were significantly more hydrophobic than the nuclear proteins quantified in P3 (Figure S4A). Hydrophobic proteins are more prone to aggregate in the brain with age^49^. Aggregated proteins are also known to be ubiquitinated^50^, and we observed a large increase in ubiquitinated proteins in P3 compared to S2 (Figure S4B). To further investigate aggregated protein accumulation in P3, we surveyed two proteomic studies^51,52^ that reported proteins enriched in the insoluble fraction in AD human brains (Figure 7E and Supplementary Table 12). We observed 37 proteins enriched in our AD P3 fraction that were also identified in one of these studies. Three of these seven proteins were APP, MAPT, and APOE, which are key players in AD pathogenesis. The other common proteins were Collagen alpha-1(XXV) chain (COL25A1), Complement C4-A, Complement Protein C3, and Transmembrane glycoprotein NMB (GPNMB) have been previously connected to AD pathogenesis. COL25A1 binds Abeta fibrils and C4-A, C3, and GPNMB support inflammation^53–56^. We observed C4-A to up-regulate in the three other fractions suggesting an increase in gene expression in addition to possible aggregation in P3. VPS29 was identified in the insoluble AD fraction in one of the studies. Thus, insoluble protein accumulation in P3 or nuclear transport dysfunction likely explains many P3/S2 discordant measurements, though not the third in which non-nuclear proteins decreased in P3 and increased in S2. These proteins were significantly enriched (FDR, 2×10^-15^) in mitochondrial proteins (Figure 7F and Supplementary Table 13). Mitochondria are dynamic organelles that can travel throughout a cell sensing local metabolic demands^57^. Furthermore, the spatial distribution of mitochondria is perturbed in AD^58^. In summary, we quantified reciprocal measurements between fractions associated with AD, which may be explained by aberrant protein trafficking.

### Quantification of AHA Degradation (QUAD)

Our quantitative MS analysis of post-mortem human brain provides a comprehensive snapshot of spatial proteoforms perturbed by AD but cannot distinguish alterations that drive pathogenesis from those that respond to it. Although not without their own limitations, AD mouse models allow studying disease progression with the bonus of employing molecular tools that are not applicable to post-mortem human brain tissue. Our human AD analysis revealed dysfunction in vesicle transport (i.e. endosomal-lysosomal network) and protein folding. Both processes can impact the kinetics of protein degradation^59,60^. This prompted us to investigate the degradation patterns in an AD mouse model with our fractionation protocol. We quantitated protein degradation in the APPswePS1delta9 AD mouse model[34] using the QUAD (Quantification of Azidohomoalanine Degradation) strategy (Figure 8A)^26^. Briefly, AD and WT mice received azidohomoalanine (AHA) in their diet for 4 days, then were returned to a normal diet for 7 days. Mice were euthanized and brains were removed after 4 days of AHA labeling (i.e. Day 0) or after 7 days on a normal diet (i.e. Day 7). A biotin-alkyne was added to the AHA inserted into the proteome via a click reaction, then the proteins were digested to peptides, and the AHA containing peptides were enriched with neutravidin beads^17^. The AHA peptides were labeled with TMT tags, then quantified with LC-MS. Protein degradation was calculated by average Day7/Day0 AHA protein ratio. The Day0 samples were pooled for each genotype within a timepoint, so the only variation in the AHA ratio was the protein degradation rate (i.e. Day7). We performed QUAD analysis on P3 and S2 fractions from AD and WT cortices at 2, 5, and 12 months. We observed five times more Abeta^42^ at 5 months compared to 2 months and more than 20-fold more Abeta^42^ at 12 months compared to 5 months consistent with previous reports (Figure S4C)^61^. We quantified 1680 unique proteins in the P3 fractions and 2991 in the S2 fraction with an overlap of 1237 quantified proteins (Figure 8B and Supplementary Table 13). There were 491 and 1105 proteins quantified in all three timepoints in the P3 and S2 fractions, respectively (Figure8 C and D). On average, there were 657 quantified proteins shared between the two fractions at each timepoint (Figure S4D-F).

**Figure 8.**
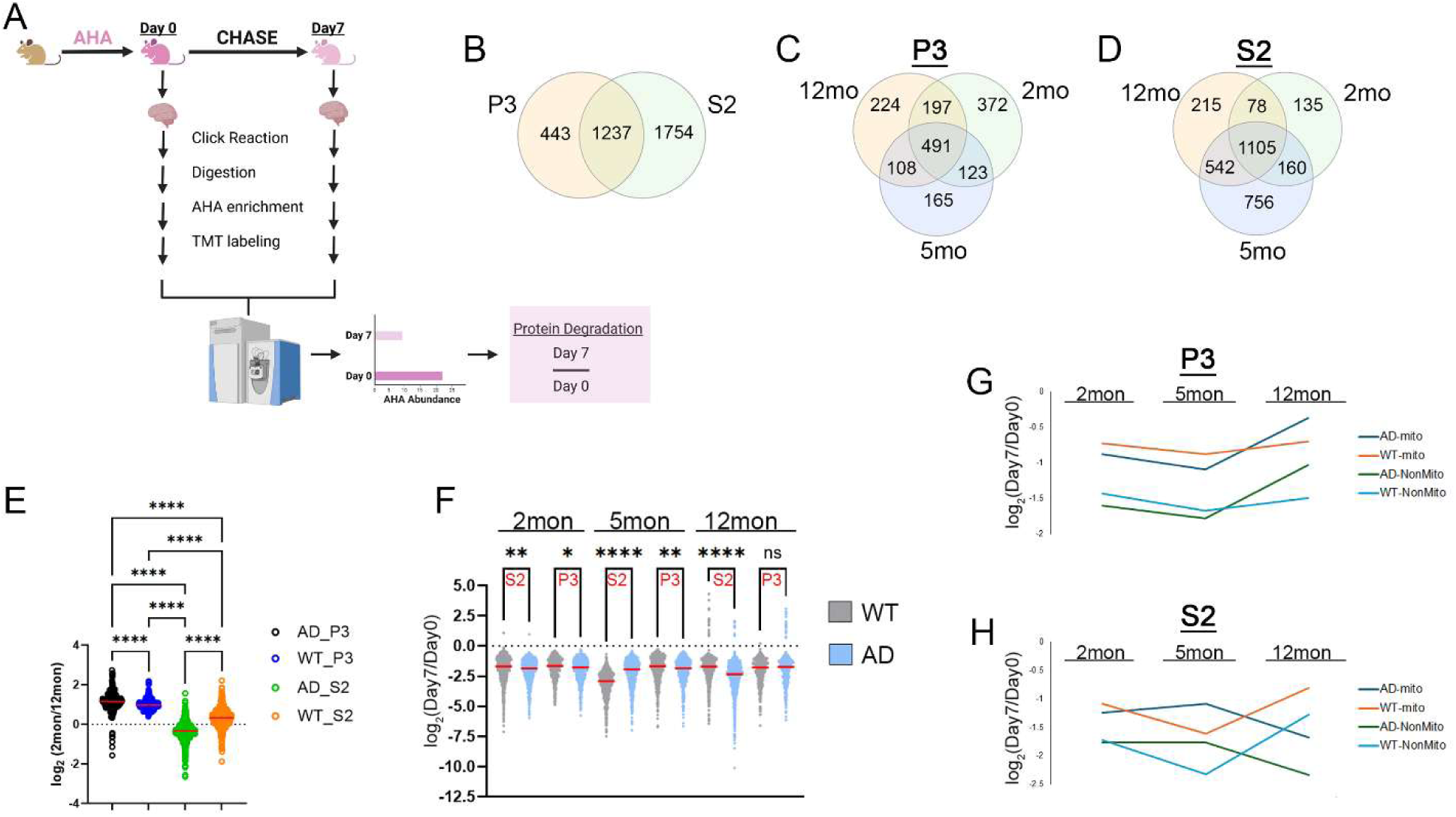
**A**, Schematic of the QUAD protocol. Venn diagrams of proteins with quantified degradations between P3 and S2 fractions (**B**), P3 at 2, 5, and 12-months(**C**), and S2 at 2, 5, and 12-months(**D**). **E**, Comparison of proteins quantified at 2 and 12months. Each circle represents a protein quantified at both 2 and 12months. The y-axis is log_2_ transformation of the ratio: 2month (Day7/Day0)/ 12month (Day7/Day0). P-values calculated by one-way ANOVA with Tukey’s multiple comparison test. **F**, Comparison of degradation rates between genotypes in each fraction and timepoint. Each circle represents the log_2_ transformation of the Day7/Day0 ratio for one protein. P-values calculated by one-way ANOVA with Tukey’s multiple comparison test. Red bars represents the medians. The mitochondrial and non-mitochondrial proteins were separated from F and their median values were plotted for P3 (**G**) and S2(**H**). Only proteins with <50% CV were plotted. P-values = **** < 0.0001, ***<0.001, ** < 0.01, * < 0.05.

We have previously demonstrated with QUAD that proteins are more stable at 12-month compared to 2-month using unfractionated mouse cortex [3]. We observed the same trend in the P3 and S2 fractions except in the AD S2 fraction, where the proteins were on average less stable at 12 months (Figure 8E). We then compared the genotypes at the same timepoint and fraction (Figure 8F). At 2 months, proteins in AD brains degraded significantly faster than in WT brains in both fractions. In later timepoints, however, a divergence between the fractions was observed. At 5 months, the S2 WT fraction possessed faster degradation than AD S2, but this trend was reversed at 12 months. In the P3 fractions, median degradation was faster in AD compared to WT at 5 months as observed at 2 months, but there was no difference observed between the genotypes at 12 months. It has been reported that the mitochondrial proteome is more stable than the whole brain proteome^26,62^. We also observed in our fractions that mitochondrial proteins were more stable than the other quantified proteins in all fractions and timepoints (Figure 8G and H). Even though clearly more stable, the mitochondrial proteome still followed the same fraction-specific degradation trend during ageing as the non-mitochondrial proteome. In summary, we observed the same protein stability trends that we and others have reported on the unfractionated brain tissue but also demonstrate that protein degradation in AD brains deviates from WT during ageing and global protein degradation patterns differ between fractions.

### ANOVA on QUAD data

Next, we identified the proteins whose degradation was significantly different between AD and WT using ANOVA. There were 238 proteins that were more stable (i.e., slower degradation) in AD compared to WT and 1,323 proteins less stable (i.e., faster degradation) in AD compared to WT. In sum, there were 1,436 proteins with altered degradation, with 113 proteins altered in more than one timepoint or fraction (Figure 9A and Supplementary Table15). In both fractions, the significant differences increased with age. At 12 months, there were stark differences between the fractions with regards to the significantly altered proteins. Proteins degraded faster in AD S2 fraction compared to WT S2, whereas most proteins were more stable in AD P3 compared to WT P3 in 12 months. The proteins that were less stable in AD S2 at 12 months were enriched in vesicle transport (i.e., vesicle transport in synapses (VTS), endosomal transport, and regulation of endocytosis) and metabolic processes and proteins that were more stable in AD P3 at 12 months were enriched in dendritic development, RNA splicing, and VTS (Figure S4G and Supplementary Table16).One of the dramatic alterations was observed with VPS35. Almost no degradation was observed in S2 WT, but efficient degradation in AD. Interestingly, this deviation in degradation between genotypes was not observed at earlier timepoints (Figure 9B).

**Figure 9.**
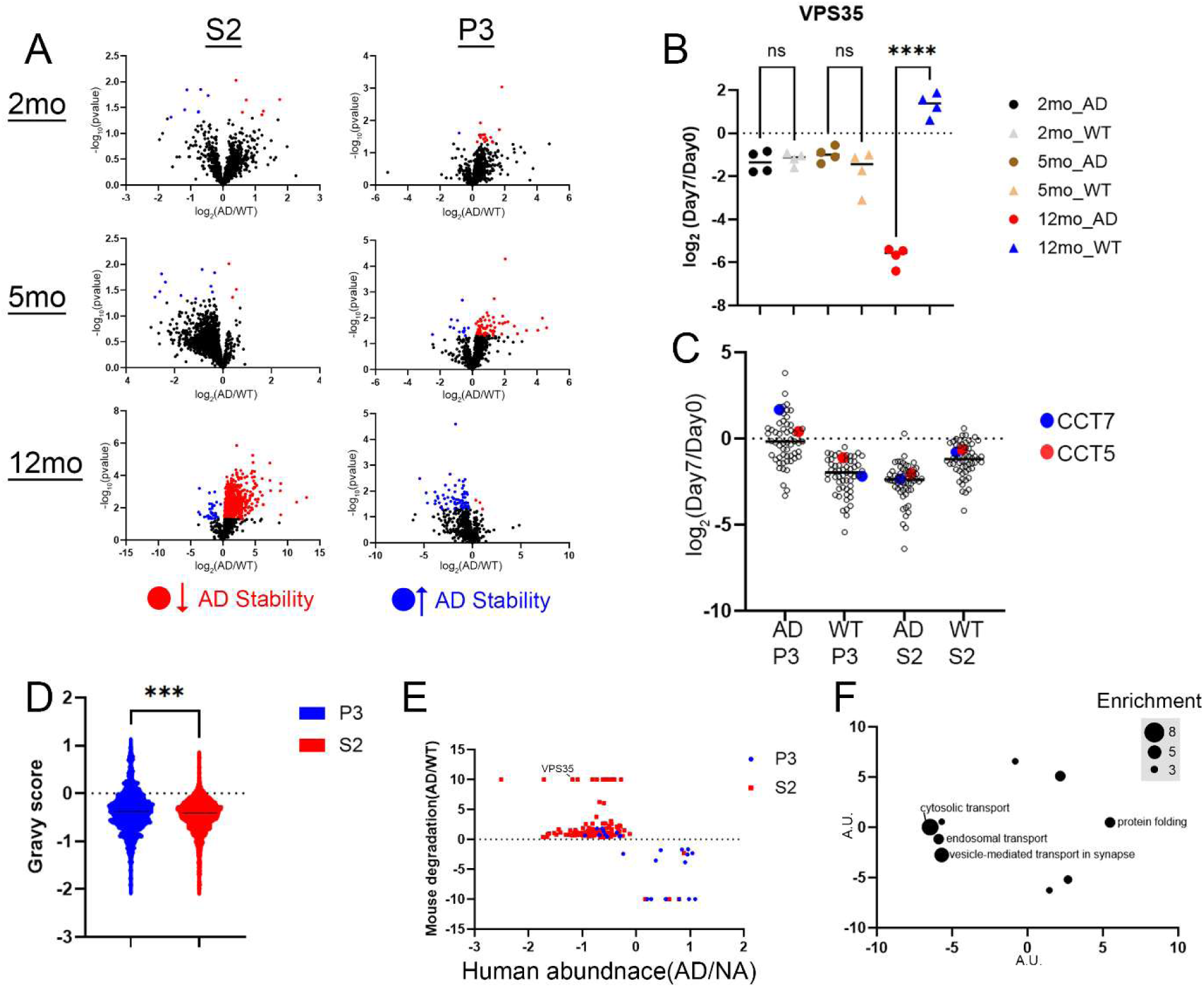
**A**, Volcano plots of the ANOVA QUAD TMT results between AD and WT brains at 2, 5, and 12months for the S2 and P3 fractions. Significant (p < 0.05) proteins are blue and red. Blue indicates that a protein degraded less in AD brains compared to WT and red indicates that a protein degraded more in AD brains compared to WT. Black circles are proteins that had a p >0.05. Y-axis is the -log_10_ of the p-value, and X-axis is the average log_2_ (AD/WT) protein Day7/Day0 values. **B**, QUAD data for VPS35. Y-axis is the log_2_ transformation of the Day7/Day0 protein ratios. Each circle represents of biological replicate. **C**, Proteins altered in both S2 and P3 at 12 months exhibiting opposite AD stability changes. The same proteins are plotted for the values in AD-P3, WT-P3, AD-S2, and WT-S2 at 12 months. Y-axis is the log_2_ transformation of the average Day7/Day0 protein ratios **D**, The gravy score (y-axis) for the unique QUAD quantified proteins in S2(red) and P3(blue). **E**, Proteins significantly different between AD and WT QUAD analysis at 12month and also significantly different in the human data in either S2(red) or P3(blue) fractions. Y-axis is the QUAD data with the average log_2_ (AD/WT) protein Day7/Day0 values and X-axis is the human data with the average log_2_ (AD/NA) protein values. **F**, GO biological processes significantly enriched in proteins that correlate between the human and mouse datasets. Circle size represents extent of enrichment. The similar GO terms are spatially closer. Axes are arbitrary units. P-values = **** < 0.0001, ***<0.001, ** < 0.01, * < 0.05.

There were 113 proteins altered in more than one timepoint or fraction. Sixty-five percent were quantified with opposite stability trends. Some of these opposite measurements were observed across different time points, which may relate to how the brain responds to different amounts of Abeta at different ages. However, we were more interested in opposite measurements occurring within the same timepoint but across different fractions, as this pattern indicates differential degradation of spatial proteoforms. There were 56 proteins altered in both S2 and P3 at 12 months exhibiting opposite AD stability changes (Figure9C). These proteins were more stable in AD P3 but less stable in AD S2 compared to WT P3 and S2 respectively. This could represent different degradation patterns between fractions as some of these proteins are known to exist in multiple subcellular compartments. For example, CAND1 (Cullin-associated NEDD8-dissociated protein 1) and CTNNB1 (beta-catenin) are localized in both the cytosol and nucleus ^63,64^. In addition, we observed discord measurement between fractions of CCT5 and CCT7 subunits of the chaperonin complex that was also perturbed in the human hippocampi analysis. Alternatively, this could represent misfolding or aggregating proteins accumulating in P3 as we observed in our human data. Like the human data, we also observed that proteins in P3 were more hydrophobic which increases the tendency of protein aggregation (Figure 9D).

### Comparison of human and mouse data

We compared the measurements that were significantly altered in the QUAD with the human hippocampal analysis. Analyzing the P3 and S2 fractions, the largest overlap with human data was the 12-month QUAD timepoint which is consistent with this being the most analogous timepoint with the human brains in terms of age and disease progression. We postulated that if a protein degrades faster in the AD mice compared to WT this would correspond to a decrease in the protein abundance in AD compared to NA, whereas if a protein degrades slower in AD mice compared to WT this would correspond to an increase in the protein abundance in AD compared to NA human samples. Thus, the correlated proteins should mathematically have opposite trends in the mouse and human analyses. Using these stipulations, we identified 155 protein measurements (27 in the P3 fraction and 128 in the S2 fraction) that correlated (Figure 9E and Supplementary Table17). For example, VPS35 decreased in human AD S2 compared to NA S2 but VPS35 degraded faster in mouse AD S2 compared to WT S2. Globally, the human-mouse correlated proteins were enriched for vesicle transport and protein folding biological pathways (Figure 9E and Supplementary Table18). We were also interested in alterations observed in human data and in the pre-pathological mouse timepoint of 2 months, because these shared proteins could be an early proteomic perturbation that triggers AD pathology. There was an overlap of six proteins (BIN1, NSFL1C, HSPA12A, CACNA2D3, DNAJA4, and PHYHIPL). Besides PHYHIPL, whose function is unknown, these proteins are linked to known AD pathogenesis hubs, including vesicle transport, protein misfolding, and calcium homeostasis. BIN1 is an AD risk factor that regulates membrane curvature, endocytosis and protein trafficking^65–67^. Similarly, NSFL1C(aka cofactor p47) regulates membrane fusion which in turn regulates autophagy, endosomal transport, and amyloid aggregation ^68,69^. CACNA2D3 is a regulatory subunit of voltage-gated calcium channels and involved in synaptic formation^70,71^. Gene variation of CACNA2D3 is a potential AD biomarker^72–74^. HSPA12A is a molecular chaperone of the HSP70 family and can bind and modulate the AD risk factor SORL1^75^. DNAJA4 is also a molecular chaperone and controls proteasomal degradation^76^. This striking concordance between QUAD and human datasets suggests a subset of AD perturbations in human tissue involved in vesicle transport and protein folding are produced by alterations in protein degradation.

## Discussion

Proteins exist as spatial proteoforms with multiple subcellular localizations. Each spatial proteoform interacts with a unique subcellular environment with access to different binding partners and metabolites (i.e. calcium). Spatial proteoform diversity is greatest in the brain, where neurons are highly compartmentalized into axonal, dendritic, synaptic, and somatic subdomains. With the analysis of brain tissue, spatial proteoforms may also be expressed in different cell types. Although this is well known, spatial proteoforms are rarely investigated with unbiased proteomic studies, most likely due to the increased analysis time. We used a simple traditional fractionation method to our advantage to ensure high overlap of spatial proteoforms among different biological fractions. Since LC-MS rarely achieves 100% sequence coverage of proteins from a complex mixture, it is possible that a protein quantified in two different fractions could be different splice isoforms or maybe the identical protein with different post-translation modifications. Endosomal pathway dysfunction is a preclinical AD hallmark, and this would suggest that there would be a large-scale disruption of protein localization^77,78^. Consistently, we observed an enrichment of vesicle trafficking and endo-lysosomal pathways (ELP) altered in AD in human hippocampi compared to NA. ELP dysfunction has been described in AD pathogenesis and predicted to be one of the earliest perturbations even before Abeta and Tau accumulation^79^. In addition, we observed alterations in other pathways previously implicated in AD including protein folding, energy production, and inflammation. Importantly, these altered pathways were observed in both a clustering-based analysis (i.e. WGNCA) and ANOVA. Because 80% of the NA group was AsymAD, our findings likely reflect proteomic changes underlying AD dementia rather than mechanisms triggered by Abeta plaques or NFT.

Our study revealed novel protein changes obscured by traditional studies and provided new spatial information to previously published AD protein alterations. We observed few proteins that were altered in all fractions indicating that most proteins are not altered simply by transcription. Alternatively, single-fraction alterations could reflect transcriptional changes for proteins targeted to only one compartment, though this is unlikely given that most altered proteins were quantified in more than one fraction. Surprisingly, most AD perturbations were restricted to one fraction. There is no doubt that AD pathology does induce transcriptional changes as evident by the numerous sequencing studies^80,81^, but our results suggest that local translation/degradation or post-translational (i.e. trafficking) regulation dictate the uniqueness of AD proteome. The sensitivity of spatial analysis is exquisitely highlighted by our data on APP, which is typically not altered in unfractionated LC-MS analysis unless only Abeta peptides are targeted for quantification^82^. APP was quantified in all fractions but only observed to be altered in P3. P3 was unique in that it contained more quantified Abeta peptides than non-Abeta peptides. Although numerous studies have analyzed one specific AD sub-proteome (i.e. insoluble fraction)^51^, we analyzed all fractions to enable us to compare perturbations between compartments as we highlighted with the discord measurements between the S2 and P3 fractions. P3 was unique due to its nuclear protein enrichment and annotated nuclear proteins were observed to possess reciprocal perturbations between AD and NA. In human AD brains, NFT accumulates on the outer surface of the nuclear envelope resulting in nuclear irregularities and NCT disruption^83–85^. This NCT dysfunction was proposed to involve the interaction between Tau and nucleoporin NUP98^86^. We observed that Tau is increased in AD P3, and also an increase in NUP160 and NUP205 in AD P3. Numerous RAN GTP binding proteins were increased in AD P3 and decreased in AD S2 consistent with a global disruption of NCT. We also observed non-nuclear P3/S2 discord proteins decreased in AD S2 and increased in AD P3 which would not be a result of NCT dysfunction. Using a VPS35 antibody, we confirmed that its S2/P3 abundance pattern is distinctly different between NA and AD. We also demonstrated that the expression in total unfractionated hippocampi is unchanged, indicating that this alteration would not been observed in unfractionated LC-MS analysis. Consistently, a meta-analysis of seven published LC-MS datasets on human unfractionated brains failed to observe a change in VPS35^27^. VPS35 is a component of the retromer complex, which transports proteins from endosomes to the trans-Golgi network or the plasma membrane. Manipulation of the retromer can disrupt protein degradation through the ELP^87^. Our quantification of a VPS35 interactor, VPS29, with the same AD P3/S2 pattern suggests a mislocalization of the entire retromer complex. The retromer complex has been observed to dysregulated in AD, and it has been suggested to be a key player in ELP dysfunction observed in preclinical AD^47^. We speculate that non-nuclear proteins enriched in P3 and diminished in S2 are aggregating or becoming less soluble causing them to shift to P3. Alternatively, these proteins could be binding insoluble proteins to prevent aggregation. Consistent with this hypothesis did observe that a portion of these proteins were previously reported to aggregate in AD brain and possessed increased hydrophobicity indicating a higher propensity to aggregation or misfolding^49^. This P3 enrichment suggests that AD brain is unable to efficiently remove insoluble protein aggregates compared to a NA brain. Interestingly, a third group of P3/S2 discord proteins was observed with decrease in P3 and increase in S2. These proteins were enriched in mitochondrial proteins, which included the AD risk factor, COX7C^88^. Mitochondria are dynamic organelles that travel to distal neuronal sites to supply ATP but are retrogradely transported back to the soma for recycling and clearance of damaged mitochondria (i.e. mitophagy)^89^. Mitochondrial dysfunction is an early event in AD pathogenesis^90^. The clearance of damaged mitochondria is impaired in AD neurons, which is due to impaired lysosome formation and autophagic flux^91,92^. We also observed an enrichment of lysosomes in S2. We hypothesized damaged mitochondria accumulate in S2 as they are not being cleared properly leading to further toxicity. Damaged mitochondria are unable to provide essential cellular activities, such as generation ATP or calcium homeostasis. Furthermore, damaged mitochondria generate ROS and if not cleared efficiently, ROS can damage protein and lipids and result in apoptosis and neurodegeneration.

Although there are many biological mechanisms proposed to cause AD dementia, our multiple unbiased bioinformatic analyses repeatedly highlighted components of the ELP and protein folding networks (PFN) as the most perturbed between AD and ND hippocampi. These two networks are intimately linked to protein degradation with the ELP clearing protein aggregates and the PFN preventing protein aggregate formation. To investigate the ELP and PFN in disease progression, we quantified protein degradation in the P3 and S2 fractions at 2, 5 and 12 months in an AD mouse model. Many reports using various analytical methods, including QUAD, have demonstrated that degradation slows with age in the unfractionated brain tissue^26^. We report that this also occurs in P3 and S2 fractions in WT brains comparing 2- and 12-month timepoints. In the AD mouse model, this trend was also observed in P3 but not S2. Examining three ages, our data suggests this decrease in degradation is not linear but there is an acceleration or plateau of degradation at 5 months followed by a decrease observed at 12 months. Additional studies are needed in more time points especially in older mice to understand the complete degradation pattern with age. AD S2 at 5 months had the largest deviation from the WT indicating that the initial increase in Abeta production has already had a profound impact on degradation. Surprisingly, we observed the reverse pattern in 12-month AD S2 with increased degradation compared to WT. There are two possible explanations for our data, which are not mutually exclusive. First, evidence suggests that degradation occurs at multiple subcellular sites. The two main degradative organelles, i.e. proteasome and lysosomes, are located throughout a cell. In neurons, they are localized to soma, nuclei, axons, dendrites, and synapses^93,94^. Furthermore, both proteasomal and lysosomes possess differential functional characteristics within different subcellular fractions, and the motility and function of these organelles can change under different conditions^95–99^. Thus, our data suggests that Abeta could alter protein degradation differentially in different subcellular compartments. Alternatively, we may not be observing changes in degradation but changes in protein localization. Like our human brain analysis, we observed alterations in our AD mouse model in the ELP which can disrupt autophagy and increase misfolded and aggregated proteins. Insoluble misfolded proteins or aggregates could exit the S2 fraction and enter P3 during centrifugation in the fractionation process. This was evident in the accumulation of known insoluble AD aggregates in our human P3 data. Thus, the quantitation of proteins with decreased stability in AD S2 and increased stability in AD P3 could be explained by a protein becoming increasingly insoluble during the seven-day chase period via misfolding or aggregation.

The AD APPswePS1delta9 mouse model is commonly employed to study AD pathogenesis, Abeta accumulation, and memory loss but has disadvantages. Most importantly, it lacks tau pathology. Although the molecular contributions of Abeta and Tau to dementia are not clear, accumulation of Abeta does occur before Tau accumulation suggesting that it is a trigger for pathogenesis. Tau accumulation, however, correlates more strongly with dementia severity than Aβ does.^100^. Therefore, we expect the intersect of our mouse and human datasets to reveal the molecular mechanisms of Abeta that contribute to dementia. For example, our human analysis revealed a dysfunction in NCT but was not detected in the QUAD dataset. NCT dysfunction is caused by hyperphosphorylated Tau^45^. Our QUAD/human intersect was enriched in ELP and protein chaperones networks suggesting these can be triggered by Abeta by itself. Although we observed few changes in 2-month AD mice, they were relevant to AD pathology. These changes may also occur in early human AD pathogenesis. One of the proteins is Bin1, which is an AD risk factor. It is unclear how BIN1 is involved in AD pathogenesis^65–67,101,102^. Most reports have suggested that it is involved in Tau pathology, but we demonstrate changes in Bin1 can be triggered in an AD model without Tau pathology.

In summary, this study introduces a spatial proteomic framework for AD that reveals a layer of biological complexity invisible to conventional unfractionated analyses. By coupling subcellular fractionation with TMT-LC-MS in human hippocampi and pairing it with QUAD across three disease stages in an AD mouse model, we demonstrate that the AD proteome is not simply defined by which proteins change, but *where* they change. Despite most AD proteins being detectable across multiple subcellular fractions, 78% of significant protein alterations are fraction specific. This finding reframes our understanding of how subcellular localization drives disease vulnerability and establishes spatial proteoforms as essential units for pathogenic analysis. The identification of retromer complex mislocalization exemplifies how spatially resolved proteomics can rescue disease-relevant signals from the noise of bulk tissue analysis. Furthermore, the detection of overlapping perturbations in BIN1, NSFL1C, HSPA12A, CACNA2D3, and DNAJA4 at the earliest pre-pathological mouse timepoint suggests that disruptions to vesicle transport fidelity and proteostasis capacity represent initiating, rather than consequential, features of AD progression. Future studies employing cell-type-specific spatial proteomics and an expanded mouse model timepoints with Tau pathology will be essential to translate these findings into mechanistic interventions and early diagnostic strategies. Together, these data demonstrate that a spatially resolved view of the proteome is indispensable for decoding the molecular architecture of Alzheimer’s disease.

## Supplementary Figures

**Supplementary Figure 1.**
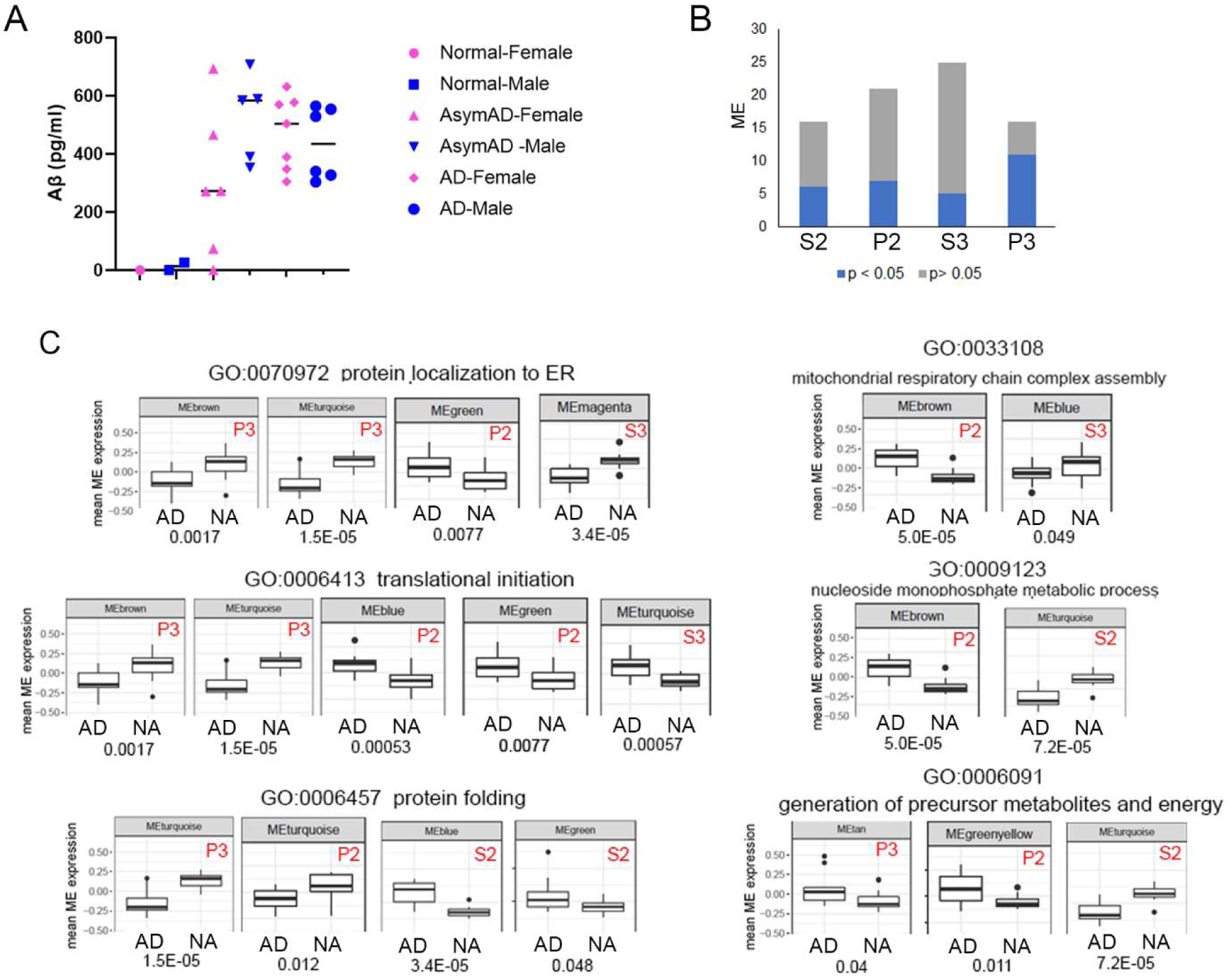
**A**, Abeta^42^ peptide concentration was calculated by an elisa on the total homogenate from the 27 hippocampi samples analyzed in this study. Y-axis is pg/ml of Abeta^42^. **B**, The number of significant and non-significant ME for each fraction as shown in Figure 2E. C, The identical GO terms from Figure 3A that were observed in different fractions with conflicting expression trends. The graphs show the mean ME expression obtained from the WGCNA. The p-value for each AD vs ND ME comparison is below each graph.

**Supplementary Figure 2.**
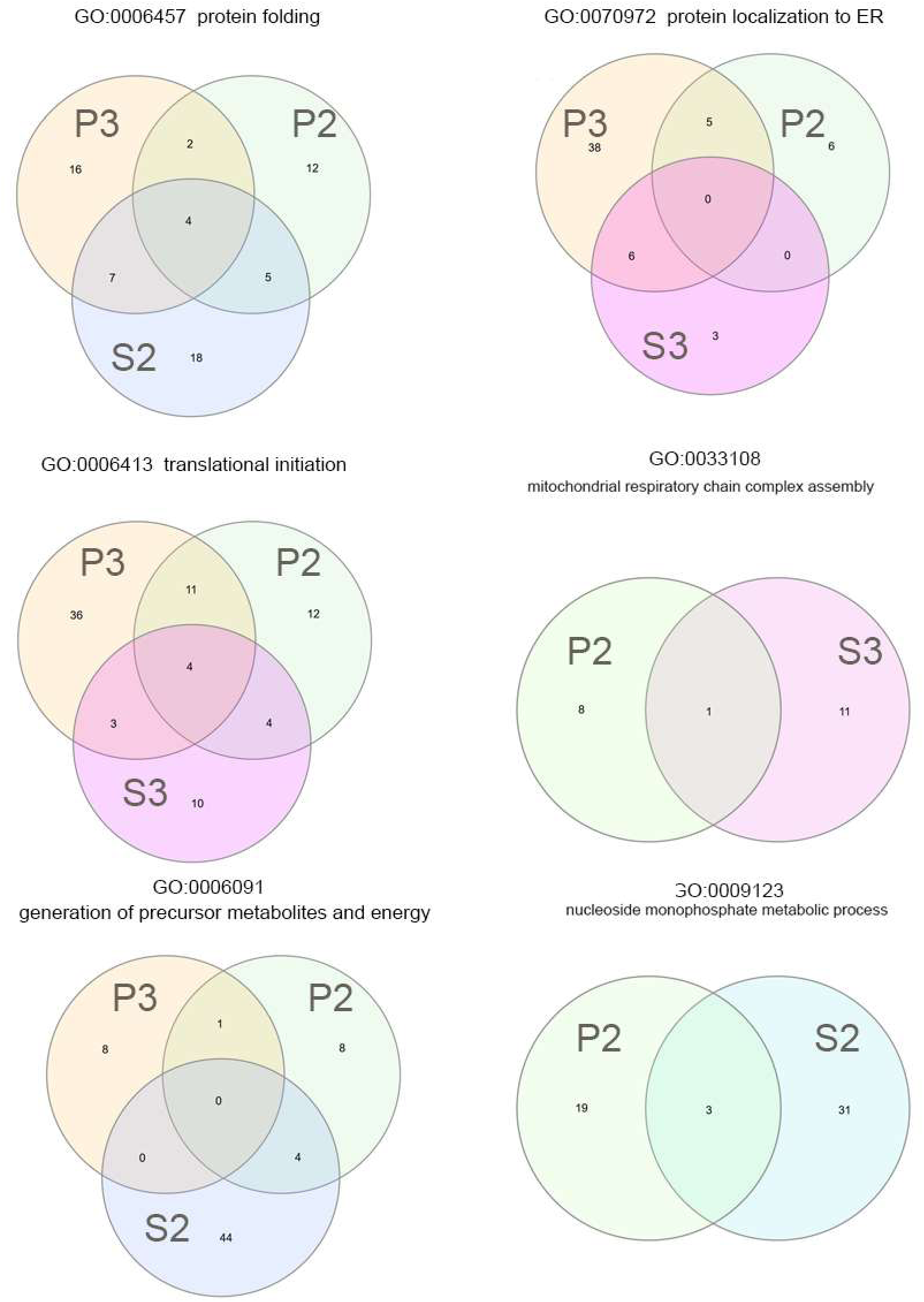
The Venn diagrams show the overlap of quantified proteins annotated to the same biological process from different fractions with conflicting expression patterns as shown in Supplementary Figure 1.

**Supplementary Figure 3.**
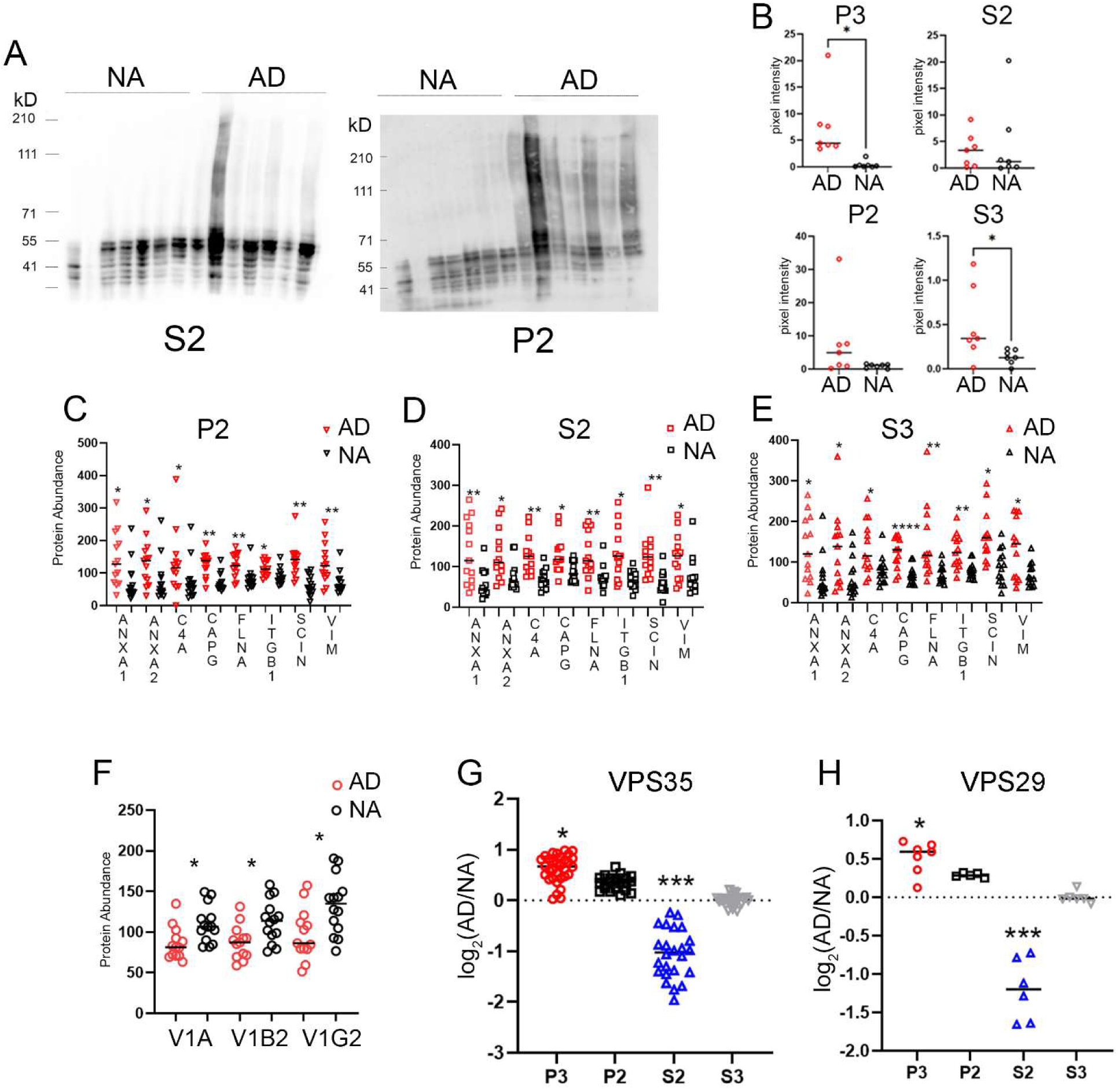
**A**, Immunoblots were probed with Tau antibody for P2 and S2 fractions. **B**, The Tau immunoreactivity from the Tau blots is plotted after being normalized to a total protein gel stain. The y-axis is normalized pixel intensity (arbitrary units). Proteins that were observed to be altered in all four fractions. Graph shows the TMT protein quantification values for the AD (red) and NA (black) samples from the P2 (**C**), S2 (**D**), and S3 (**E**) fractions. **F**, The TMT protein quantification values for proteins (ATP6V1A, ATP6V1B2, ATP6VG2) of the V-ATPase complex in the P3 fraction. The quantified peptides for VPS35 (**G**) and VPS29 (**H**) that were up-regulated in the P3 fraction (red) and down-regulated in the S2 fraction (blue). The y-axis is the log^2^ of ratio of the AD peptide measurement over the NA peptide measurement. p-values: * < 0.05, ** < 0.01, ***< 0.005, **** < 0.0001

**Supplementary Figure 4.**
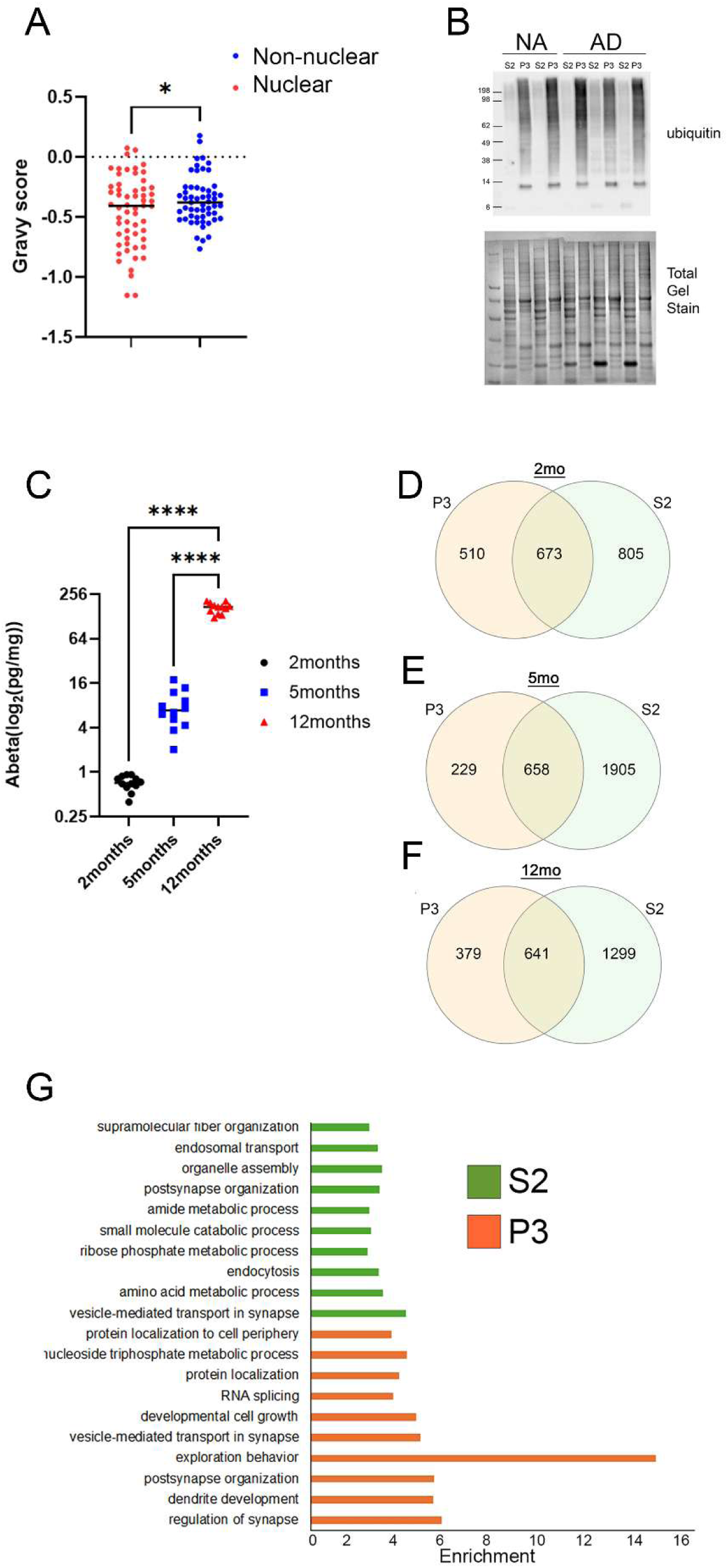
**A**, Hydrophobicity of nuclear and non-nuclear proteins quantified in the P3 fraction. **B**, Ubiquitin immunoreactivity on an immunoblot(upper panel) comparing S2 and P3 fractions. Lower panel is total protein stain. **C**, Abeta^42^ peptide concentration was calculated by an elisa on the total homogenate from AD mice used in this study. Y-axis is log^2^ pg/ml of Abeta^42^. The overlap of quantified AHA proteins between the S2 and P3 fractions at 2months(**D**), 5months (**E**), and 12months (**F**). **G**, Significant (pvalue < 0.0003) enriched GO biological processes for S2AD12 (S2-green) and P3AD12(P3-orange) using a 5% FDR filter.

## Supplementary Information

**Supplementary Table 1.** Meta data for the post-mortem brain tissue including age, gender, and pathology report.

**Supplementary Table 2**. All the proteins identified by at least two peptides of different sequences are listed by their Uniprot accession number and the fraction where they were observed. Data plotted in Figure 2A, B, and C.

**Supplementary Table 3.** The significantly enriched GO cellular compartments using a 5% FDR filter for each fraction. Data represented in Figure 2D.

**Supplementary Table 4.** The module eigenprotein (ME) calculated by MetaNetwork WGNCA and p-values for differences between AD and ND samples. Data represented in Figure 2E.

**Supplementary Table 5.** The enriched GO biological processes for the ME altered in AD. Data depicted in Figure 3A.

**Supplementary Table 6.** Proteins observed to be significantly altered in the volcano plots in Figure 4A.

**Supplementary Table 7.** Comparison of ANOVA with a previously published meta-analysis of seven proteomic TMT datasets on human unfractionated AD non-hippocampal brain tissue.

**Supplementary Table 8.** The significantly enriched GO biological processes from the ANOVA as shown in Figure 4D.

**Supplementary Table 9.** Proteins that correlated with Tau abundance.

**Supplementary Table 10.** The significantly enriched GO biological processes that were enriched in shared proteins that exhibited similar expression patterns in the P2 and P3 fractions.

**Supplementary Table 11.** The significantly enriched GO molecular processes that were enriched in discord proteins from the S2 and P3 fractions.

**Supplementary Table 12.** Proteins upregulated in P3 AD that were previously reported in the AD insoluble fraction as depicted in Figure 7E.

**Supplementary Table 13.** Proteins depicted in Figure 7F.

**Supplementary Table 14**. All the proteins quantified with their Day7/Day0 protein ratios in P3 and S2 fractions at 2,5, and 12months. The data for each fraction and timepoint is displayed on a separate worksheet. Mitochondrial proteins are marked that were plotted in Figure 8G and H.

**Supplementary Table 15**. All the QUAD quantified proteins that were significantly (p < 0.05) different between AD and WT as plotted in Figure 9A.

**Supplementary Table 16**. The GO biological process terms enriched in the proteins that were less stable in AD S2 at 12months and proteins that were more stable in AD P3 at 12months as depicted.

**Supplementary Table 17.** Proteins quantified in the S2 and P3 fractions in both human and mouse brains.

**Supplementary Table 18**. The GO biological process terms enriched in the proteins that quantified in the S2 and P3 fractions in both human brains and mouse brains at 12month old depicted in Supplementary Figure 9F.

## Notes

### Competing Interest Statement

The authors have declared no competing interest.

### Funding

- NIH Common Fund: 1R01 AG077046-01

